# Generation of human hindlimb/genital tubercle progenitors from pluripotent stem cells

**DOI:** 10.64898/2026.04.14.718471

**Authors:** Sude Uyulgan, Sofia Sedas Perez, Kavitha Chinnaiya, Matthew Towers, Anestis Tsakiridis

**Affiliations:** Centre for Stem Cell Biology, School of Biosciences, The University of Sheffield, Sheffield, UK; Neuroscience Institute, The University of Sheffield, Sheffield, UK; School of Biosciences, University of Sheffield, Western Bank, Sheffield, UK

## Abstract

The hindlimbs and genital tubercle arise from a bipotent posterior lateral plate mesoderm (pLPM) progenitor population and share common genetic programmes, reflecting their deep evolutionary homology. Perturbations of this shared developmental programme underlie several congenital conditions, yet most insights into the divergence of pLPM fates come from traditional model organisms, emphasising the need for a human-based system. Here, we report the WNT-FGF-BMP-dependent differentiation of human pluripotent stem cells into pLPM-derived hindlimb/genital tubercle progenitors (HGTps). BMP signalling drives transcriptomic changes reminiscent of the trunk-to-tail transition—a major reorganisation of the embryonic body plan that initiates pLPM/HGTp specification. We further show that retinoic acid acts biphasically: early exposure blocks these changes, whereas late exposure refines HGTp fate by suppressing alternative pLPM derivatives. Single-cell profiling reveals a spectrum of uncommitted progenitors and cells acquiring genital tubercle or hindlimb character, which map to posterior appendage populations of Carnegie stage 12–18 human embryos. Differentiating mesenchyme self-organises alongside a surrounding epithelium, mirroring appendage development *in vivo*. Grafted HGTp-derived spheroids incorporate into the genital tubercle-forming region of chick embryos, revealing their positional competence. Collectively, our work establishes a platform for the reverse engineering and disease modelling of human HGTps.

**Blurb:** Hindlimbs and external genitalia share a common developmental origin. We establish a human stem cell model to investigate their specification in vitro.

## Introduction

The hindlimbs and genital tubercle—the precursor of the external genitalia—form during the trunk-to-tail transition, which redirects lateral plate mesoderm (LPM) production from anterior (aLPM) to posterior (pLPM) regions of the embryo [1–3]. During this transition, bi-fated neuromesodermal progenitors (NMPs), which give rise to both spinal cord neurectoderm and paraxial mesoderm, relocate from the node/primitive streak border to the chordoneural hinge, where they contribute to the tailbud [4]. Another key event that occurs during the trunk-to-tail transition is the elimination of Retinoic Acid (RA) signalling from the late primitive streak and early tailbud, which along with other signals including FGF and WNT, specifies regional identity along the antero-posterior (A-P) axis by establishing the co-linear pattern of *Hox* gene expression [4].

Comparative and functional studies suggest that a HoxD-dependent genetic programme underlying external genital formation predates and was co-opted for the patterning of paired appendages during vertebrate evolution, representing a striking example of deep homology [5]. Supporting this relationship, the mesenchymal components of the hindlimbs and genital tubercle arise from a common progenitor pool within the pLPM, which we term hindlimb/genital tubercle progenitors (HGTps) [1–3,6,7].

Along the medio-lateral axis of the pLPM, the HGTp field segregates into medial hindlimb progenitors and lateral genital tubercle progenitors, which later converge at the ventral midline during body wall closure to form the external genitalia [7,8]. HGTp-derived pericloacal mesenchyme subsequently forms the primordium of the genital tubercle—a process dependent on instructive signals including Sonic hedgehog (Shh) produced by the adjacent endoderm-derived cloaca [9,10]. Both limb and genital tubercle mesenchyme are surrounded by an epithelium, and reciprocal interactions between these two tissue layers are essential for the patterning and growth of these appendages [11].

The shared developmental origin of the hindlimbs and genital tubercle is indicated by the co-expression and lineage tracing of genes encoding key transcription factors, including *Tbx4* and *Islet1* (*Isl1*), in mouse HGTps [1,6,12]. Notably, the transition of HGTps into distinct medio-lateral fates is marked by the specific upregulation of *Tbx5* in genital tubercle mesenchymal progenitors [7], which is also an essential regulator of forelimb development [14,15]. In humans, recent single cell RNA-sequencing (scRNA-seq) analysis of the reproductive system classifies genital tubercle mesenchymal progenitors as a *TBX4*^+^*TBX5*^+^ population [15] (**S1A Fig**).

In the mouse, Gdf11, and potentially other TGFβ/BMP family ligands acting through TGFβ receptor 1 (TGFβR1), directs differentiation of Tbx4^+^Isl1^+^ HGTps toward hindlimb (Tbx5 Isl1^−^) and genital tubercle (Tbx5 Isl1^+^) fates [1–3,16,17]. This process is tightly coupled to the trunk-to-tail transition and NMP relocation, which is also driven by Gdf11 [1,18]. The bipotency of HGTps is further exemplified in *Tgfbr1*-deficient mouse embryos, which fail to induce *Tbx5* and instead form a second pair of hindlimbs at the expense of genital structures [2].

The developmental and evolutionary relationship between hindlimbs and external genitalia is reflected clinically, as congenital conditions, such as sirenomelia (mermaid syndrome), caudal regression syndrome and VACTERL association, affect both appendages [19–21]. Human pluripotent stem cells (hPSCs) provide a powerful platform for modelling pLPM cell fate decisions and associated birth defects. Although the production of aLPM and their derivatives (e.g., cardiac mesoderm) from hPSCs has been reported [22–24], the efficient derivation of human pLPM and HGTps remains challenging. Previous attempts to generate human hindlimb-like progenitors via chemically induced reprogramming of adult somatic cells, or by differentiating hPSCs into limb-like cartilage, failed to detect *TBX4* [25,26]. As *TBX4* marks HGTps and their derivatives, the identities of the differentiated cells obtained in these studies remain ambiguous.

Here, we address the need for a human system by directing hPSCs to generate HGTps. We show that early BMP stimulation diverts differentiating hPSCs away from an NMP-like trajectory toward pLPM, and subsequently an HGTp fate. Early activation of RA signalling antagonises this process by blocking trunk-to-tail-transition-like transcriptome changes and lumbosacral *HOX* gene activation. By contrast, late RA stimulation refines HGTp specification by suppressing alternative pLPM-derived fates, such as endothelial lineages. Single cell RNA sequencing confirmed the presence of a continuum of uncommitted progenitors and cells acquiring genital tubercle or hindlimb identity in our HGTp cultures; these were found to resemble posterior appendage cell populations in Carnegie stage 12–18 human embryos. Xenografting HGTps into the chick embryo shows that they retain positional information and incorporate into the genital tubercle-forming region. Together, our work provides a tractable human system for studying hindlimb and genital tubercle development.

## Results

### BMP signalling diverts differentiating hPSCs from an NMP to a pLPM/HGTp fate

In the mouse embryo, pLPM progenitors emerge in the posterior primitive streak/caudal lateral epiblast region, which is near the NMP-containing regions [4,27–29], and exhibit high BMP activity [4,30–32]. Building on this observation, and the reported BMP-mediated developmental plasticity between NMPs and pLPM in the mouse [27,31,32], we examined how this signalling pathway influences fate choice between these progenitor populations during differentiation of hPSCs. Using an induced pluripotent stem cell (iPSC) line (113-CR-3) [33], we generated NMPs through our established 3-day protocol involving treatment with FGF2 and the WNT agonist/GSK3 inhibitor - CHIR99021 (CHIR) [34], in the presence or absence of BMP4 (**Fig 1A**). Quantitative PCR (qPCR) at day 3 (D3) showed that BMP4 treatment (20 ng/ml) led to strong induction of pLPM/HGTp markers and canonical BMP downstream targets such as *ISL1*, *HAND1* and *GATA4* (**Fig 1B**) [30,35]. Similar results were obtained with lower BMP4 concentrations (**S1B Fig)**. Immunofluorescence analysis confirmed that ISL1⁺ nascent HGTp cells represented the majority (∼60%) of D3 cultures (**Fig 1C**). BMP-driven pLPM induction was accompanied by reduced expression of the NMP/early mesoderm marker *TBXT* and increased expression of the EMT (Epithelial-to- Mesenchymal Transition)/pLPM marker *TWIST1* [30,36] (**Fig 1B**, and **1C**). These data indicate that BMP signalling promotes an EMT characteristic of pLPM progenitors.

**Fig 1.**
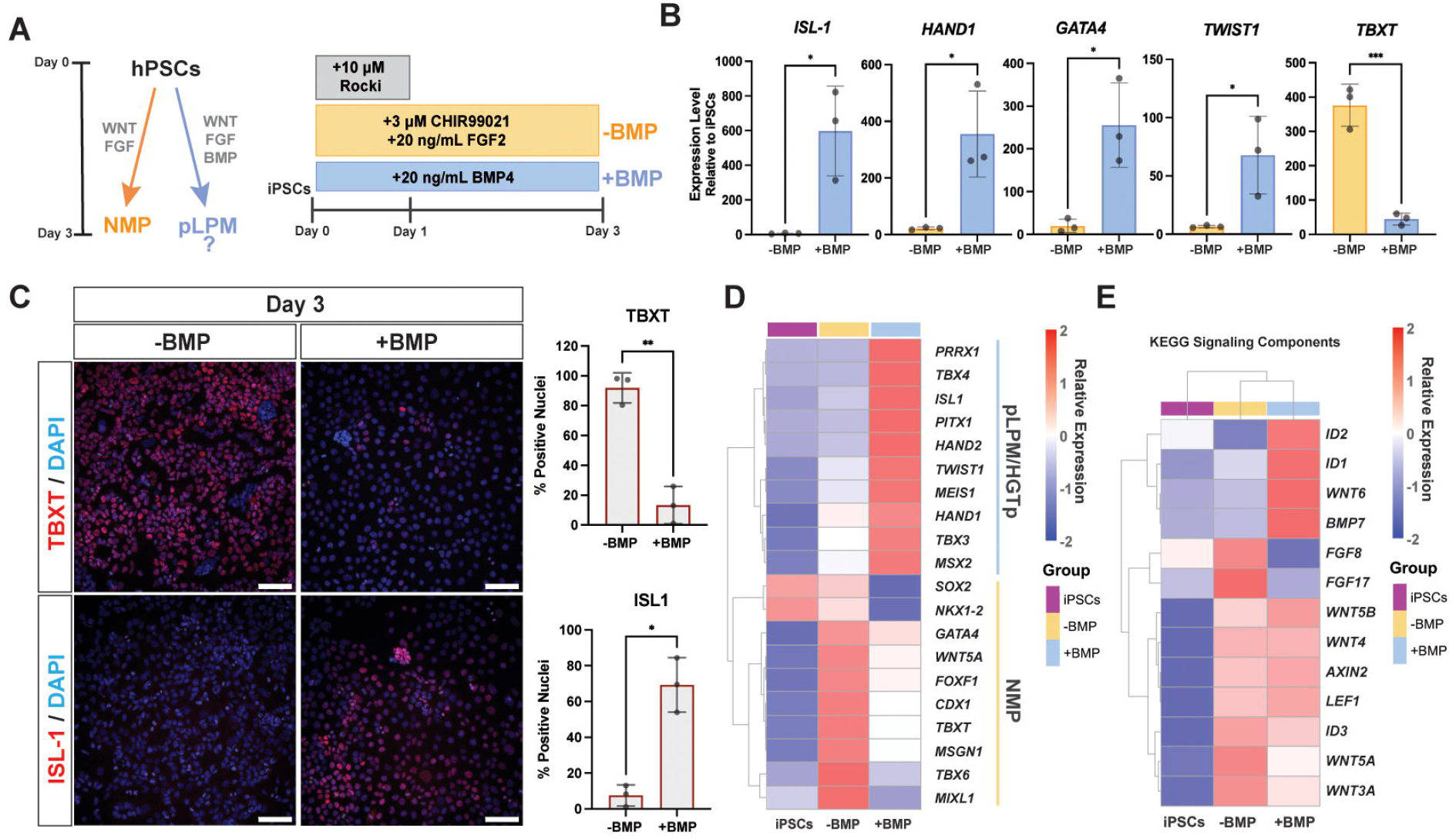
BMP signalling diverts differentiating hPSCs from an NMP to a pLPM/HGTp fate. **(A)** Treatment conditions used to induce NMPs and pLPM cells from hPSCs. **(B)** RT-qPCR- based expression analysis of key NMP and pLPM markers after 3 days of iPSC differentiation toward NMPs in the presence and absence of BMP4 (n = 3 independent experiments). Relative expression levels to iPSCs were normalised to GAPDH gene expression (* p<0.05, ** p<0.01, *** p<0.001, mean ±SD) (paired, two-tailed t-test). **(C)** Immunofluorescence analysis of the expression of TBXT and ISL1 on day 3 of differentiation. Scale bars represent 100 µm. Image analysis of the percentage of nuclei positive for TBXT and ISL1 protein expression is also shown. Graph shows mean values (n = 3 independent experiments) (* p<0.05, ** p<0.01, *** p<0.001, mean ±SD) (unpaired t-test with Welch’s correction). **(D)** Heatmap of differentially expressed genes showing key NMP and pLPM/HGTp markers from bulk RNA-seq analysis on day 3. Expression values are shown as Z-scores across samples. **(E)** Heatmap of differentially expressed genes showing KEGG signalling components on day 3. Expression values are shown as Z-scores across samples.

To further explore how BMP signalling drives pLPM specification, we performed bulk RNA sequencing (RNA-seq). We observed dramatic transcriptomic differences between D3 NMPs and their BMP-treated counterparts, with 1817 and 1660 genes significantly up- and downregulated respectively upon BMP stimulation (DESeq2; padj<0.05; |log2FC|>1)) (**S1C Fig** and **S1 Table**). In line with our qPCR/immunofluorescence data, BMP stimulation significantly induced numerous pLPM-associated/BMP target genes, including *HAND1/2* [30,37–40] and *PRRX1* [41] as well as the HGTp markers *PITX1, TBX4* and *ISL1* [1,6,42,43] (**Fig 1D**)(**S1 Table**). Conversely, transcripts marking an NMP and paraxial mesoderm identity (*TBXT, SOX2, MSGN1, TBX6*) [27,30] were markedly reduced (**Fig 1D**)(**S1 Table**). We also observed an enrichment in BMP (*ID1/2, BMP7*) and WNT (*WNT6*) signalling components that are expressed in the pLPM progenitor niche *in vivo* [30], whereas WNT/FGF-related transcripts enriched in the NMP/paraxial mesoderm progenitor-harbouring regions (e.g., *WNT3A/5B, AXIN2, FGF8*) were preferentially expressed in hPSC-derived NMPs (**Fig 1E**)(**S1 Table**).

Together, these results demonstrate that BMP signalling stimulation, combined with WNT and FGF agonists during pluripotency exit, directs differentiating hPSCs toward a pLPM/HGTp fate at the expense of an NMP fate.

### Induction of human genital tubercle progenitors

We next examined the fate of hPSC-derived D3 pLPM/HGTp cells upon further culture in the presence of WNT, FGF and BMP agonists up to D5 (**Fig 2A**). Bulk RNA-seq analysis of D0, D3 and D5 cultures revealed two major transcriptional waves: first, a cohort of pLPM– associated transcripts (e.g., *MSX1/2, HAND1, TBX3*) [14,44] was induced early and maintained from D3 to D5 (**Fig 2B**)(**S2 Table**); second, genes defining HGTps (e.g. *PITX1*, *TBX4*) alongside limb bud/genital tubercle progenitor mesenchyme markers (*IRX3*, *PRRX1, TBX5, DKK2*) [7,41,45–48] and EMT regulators (*TWIST1, SNAI1, ZEB1/2*)[36,49,50] generally showed a progressive increase in expression between D0 and D5 (**Fig 2B**)(**S2** and **S3 Tables**). The upregulation of *TBX5* by D5 strongly indicates the specific induction of genital tubercle progenitors from pLPM/HGTp cells [2,7]. This is supported by the concomitant downregulation of *TBXT* (**Fig 1B**, and **1C**), a marker of NMPs/presomitic mesoderm that also labels the allantois, the only other posterior tissue reported to express *TBX5* [51–53].

**Fig 2.**
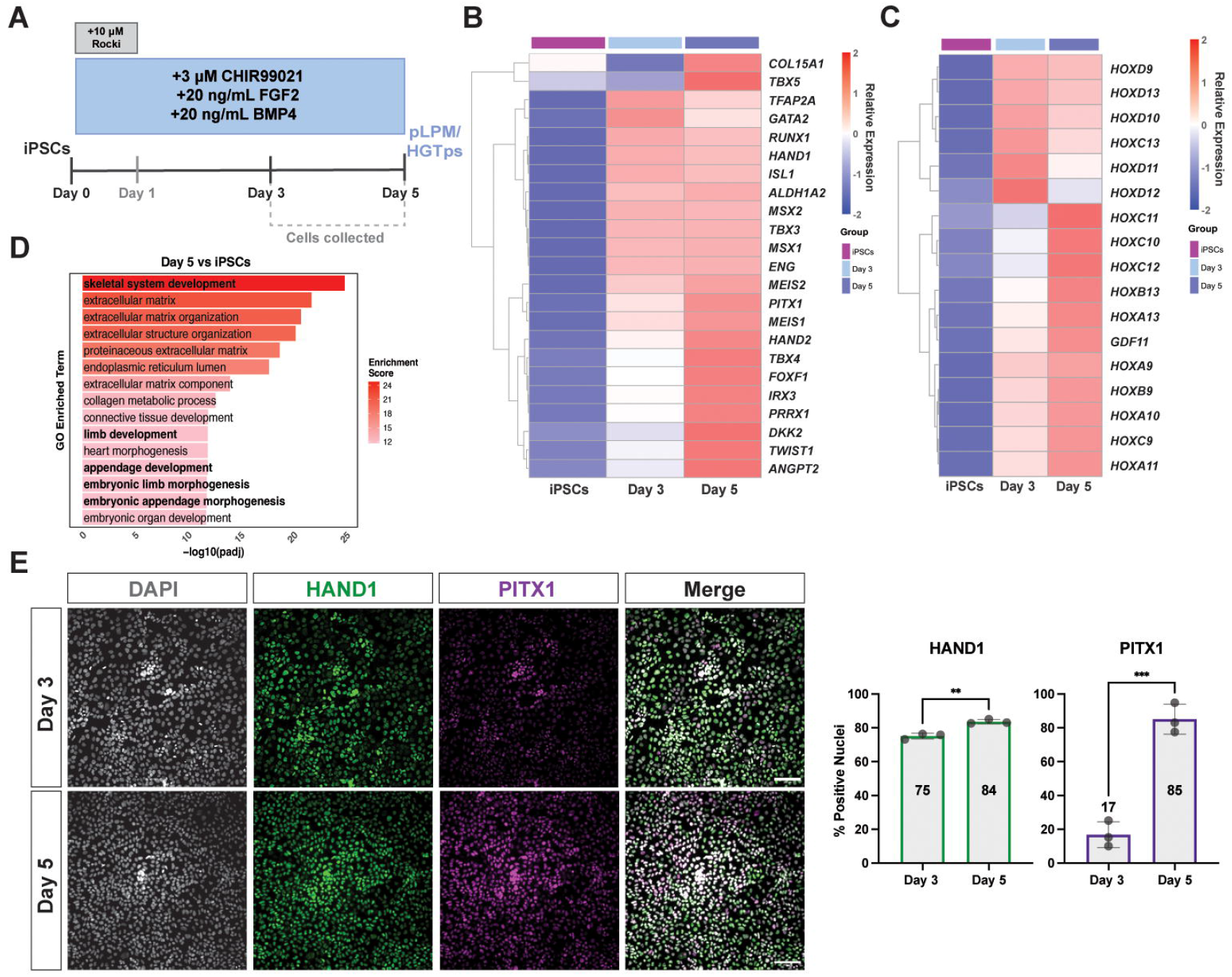
Induction of human genital tubercle progenitors. **(A)** Schematic of the treatment conditions used to induce human genital tubercle progenitors from iPSCs. **(B)** Heatmap of differentially expressed key pLPM/HGTp/genital tubercle mesenchyme markers from bulk RNA-seq on iPSCs and differentiation day 3 and day 5 cells. Expression values are shown as Z-scores across samples. **(C)** Heatmap of indicated differentially expressed genes (HOXPG9-13 members and GDF11) from bulk RNA-seq on iPSCs and differentiation day 3 and day 5 cells. Expression values are shown as Z-scores across samples. **(D)** Top 15 significantly enriched Gene Ontology (GO) terms for upregulated genes in day 5 cells vs iPSCs. **(E)** Immunofluorescence analysis of the expression of HAND1 and PITX1 on days 3, and 5 of differentiation. Scale bars represent 100 µm. Image analysis of the percentage of nuclei positive for PITX1 and HAND1 protein expression is also shown. Graph shows mean values (n = 3 independent experiments) (* p<0.05, ** p<0.01, *** p<0.001, mean ±SD (unpaired t-test with Welch’s correction).

We also detected upregulation of transcripts associated with a vascular/endothelial character (*ENG, ANGPTL2, VEGFB/C*) [54–56] consistent with the reported capacity of pLPM progenitors to generate these lineages [31,57] (**Fig 2B**)( **S2** and **S3 Tables**). Finally, combined BMP, WNT and FGF stimulation triggered upregulation of 5’ lumbosacral *HOXA-D* genes belonging to paralogous groups (PG)(9–13) and *GDF11*, indicative of trunk-to-tail–like transcriptional changes and the acquisition of a posterior fate (**Fig 2C**). Notably, the robust induction of 5′ *HOXD* PG(9–13) genes at D3 was more pronounced than that of the corresponding 5′ *HOXA–C* PG(9–13) genes (**Fig 2C**). This selective enrichment of 5′ *HOXD* expression likely reflects HGTp/genital tubercle specification [58], consistent with the absence of 5’ *HOXB* expression in this structure [59].

Gene Ontology (GO) enrichment analysis (hypergeometric test, padj < 0.05) further supported the acquisition of a strong HGTp/hindlimb/genital tubercle mesenchymal identity by D5, with top-enriched biological process terms related to skeletal, limb, cartilage, appendage and connective tissue development (**Fig 2D**)(**S4 Table**). Immunofluorescence analysis corroborated these findings at the protein level demonstrating that the proportion of cells expressing the pLPM/BMP target marker HAND1 increased modestly between D3 and D5, while the number of cells exhibiting expression of the HGTp marker PITX1 increased dramatically, rising from ∼20% at D3 to ∼85% by D5 (**Fig 2E**). Similar results were obtained using a second independent human iPSC line (SFCi55-ZsGr) [60] as well as a human embryonic stem cell line (H9/WA09) [61] (**S2 Fig**).

Together, these data indicate that sustained stimulation of BMP, WNT, and FGF signalling drives the progressive differentiation of hPSC-derived pLPM cells into HGTps and their mesenchymal derivatives, alongside an endothelial subpopulation.

### RA signalling refines HGTp specification and differentiation

Our RNA-seq analysis revealed that the BMP-WNT-FGF-driven induction of pLPM/HGTps and, subsequently, genital tubercle mesenchymal cells, coincides with progressive upregulation in the expression of the gene encoding the RA-synthesising enzyme ALDH1A2 between D3-5 (**Fig 2B**). As RA signalling is a well-established regulator of LPM patterning [62,63], and *Aldh1a2* is expressed in the genital tubercle in mouse embryos [46,64], we next sought to determine how RA signalling influences HGTp specification/differentiation.

To distinguish stage-specific roles of RA signalling, we implemented two RA treatment regimes in combination with BMP, WNT, and FGF agonists during hPSC differentiation (**Fig 3A**): RA was added (i) during pluripotency exit at D0 (“early RA”); and (ii) after pLPM/HGTp induction (marked by lumbosacral *HOX* gene expression) at D5 (“late RA”). RNA-seq analysis at D10 revealed that early RA strongly suppressed posterior identity, resulting in marked depletion of lumbosacral *HOX* PG(*10–13*) members and pLPM/HGTp-associated transcripts, including *TBX4, PITX1,* and *ISL1* (**Fig 3B-D**)(**S5 Table**). By contrast, late RA, indicated by strong upregulation of the RA target gene *CYP26A1*, led to downregulation of endothelial and vascular markers (*PDGFA, FLT1, KDR*) [65,66], while maintaining expression of HGTp/genital tubercle markers (*ISL1, PITX1, TBX4, TBX5*) and lumbosacral *HOX* genes observed under RA-free conditions **(Fig 3B–D**)(**S6, S7 Tables**). Similar results were obtained following differentiation of SFCi55-ZsGr and H9 hPSCs (**S3 Fig**).

**Fig 3.**
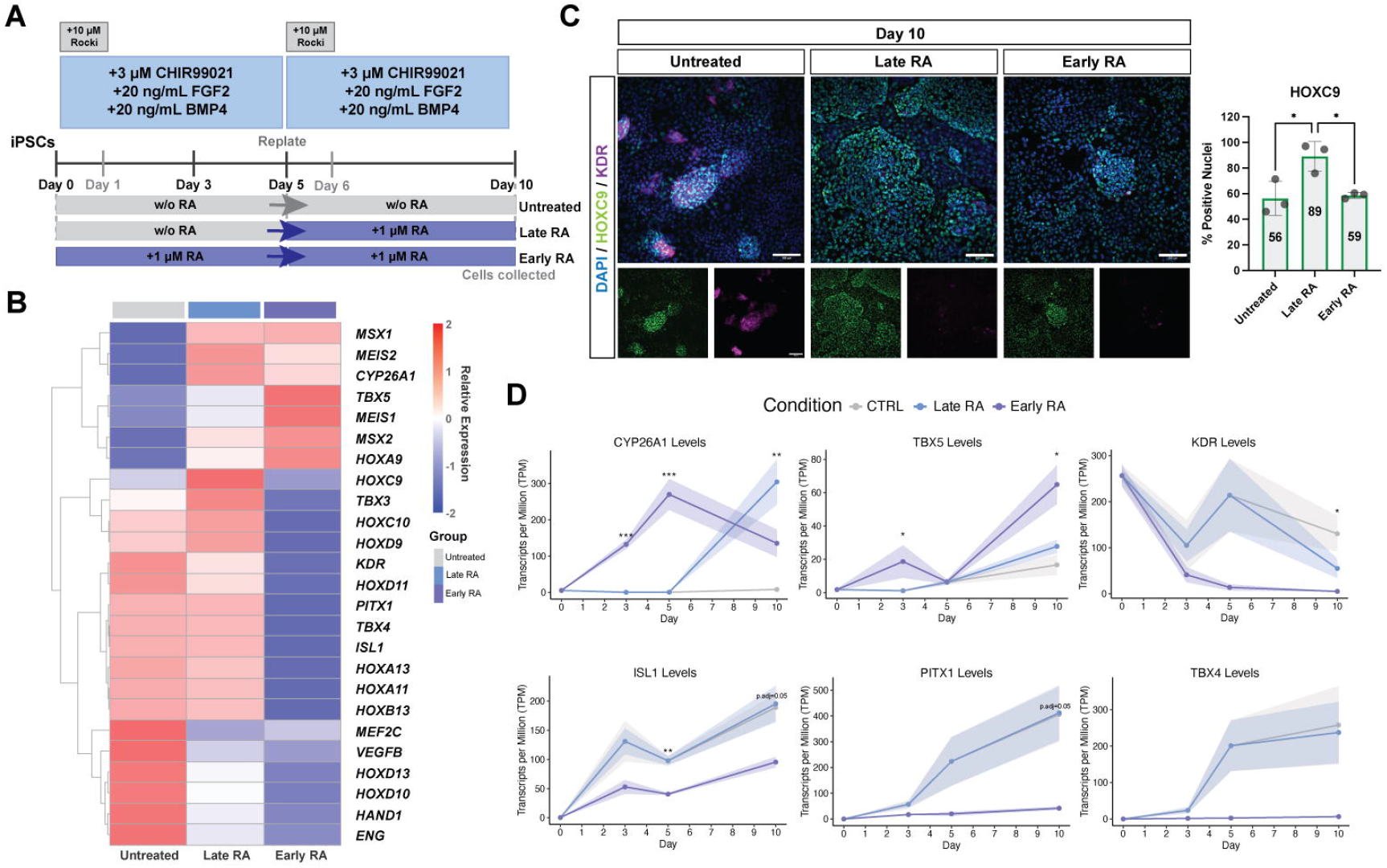
RA signalling refines HGTp specification/differentiation. **(A)** Schematic of the treatment conditions used to test different regimens of RA treatment during hPSC differentiation toward HGTps. **(B)** Heatmap of differentially expressed pLPM/HGTp, vascular/endothelial markers and posterior HOX genes from bulk RNA-seq on day 10 untreated, late RA and early RA treated cells. Expression values are shown as Z-scores across samples. **(C)** Immunofluorescence analysis of the expression of HOXC9 and KDR on day 10 untreated, late RA and early RA treated cells. Scale bars represent 100 µm. Image analysis of the percentage of nuclei positive for HOXC9 protein expression is also shown. Graph shows mean values (n = 3 independent experiments) (* p<0.05, ** p<0.01, *** p<0.001, mean ±SD) (one way ANOVA and Tukey’s multiple comparison test). **(D)** RNA-seq timecourse analysis of the expression levels of key RA target, genital tubercle mesenchyme, vascular/endothelial, pLPM/HGTp markers. TPM is calculated and analysed using one-way ANOVA and Tukey’s test to calculate adjusted p value (* adj p<0.05, ** adj p<0.01, *** adj p<0.001, mean ±SEM).

Together, these data reveal a biphasic role for RA signalling: early exposure at the exit from pluripotency blocks posterior *HOX* gene activation and trunk-to-tail transition-like transcriptome changes, whereas late exposure refines HGTp specification and further differentiation toward a genital tubercle fate by suppressing the emergence of pLPM endothelial/vascular derivatives.

### Single cell RNA-sequencing reveals the co-differentiation of HGTp derivatives and ectodermal cells

As sustained BMP–WNT–FGF signalling followed by a late RA pulse depleted off-target cell populations at D10 (**Fig 3**), we characterised the cellular composition of these cultures using droplet-based scRNA-seq. We obtained 8,292 cells that passed quality control (**S4 Fig**), which were bioinformatically allocated to three distinct clusters based on identification of well-established marker gene expression (**Fig 4A**). Cluster 1 contained most cells (77%), which mainly exhibited a HGTp character, marked by the simultaneous presence of HGTp (e.g., *TBX4, PITX1, ISL1*) and the genital tubercle mesenchyme-related transcript *TBX5,* together with *HOX* PG (9-13) genes (**Fig 4B-E**; **S8 Table**). This mixed identity fraction included presumptive *TBX4^+^ISL1^+^*HGTp progenitors (33%) and *TBX5^+^*-expressing subpopulations including *TBX4^+^TBX5^+^ISL1^+^* (16%), and *TBX5^+^ISL1^+^*(5%), representing gradual commitment toward a genital tubercle mesenchyme fate (**S5A-B Fig)**. A minor subpopulation *HOXB13*-expressing cells, likely representing uncommitted tailbud mesoderm derivatives, was also detected (**S5C Fig**). The small number of *HOX* PG(4-5)*^+^TBX5^+^*(4-6% in cluster 1) cells indicates minimal presence of forelimb/aLPM derivatives (**S5D Fig**) consistent with the near absence of markers of a definitive cardiac differentiation programme (**S6 Fig**). Similarly, we could not detect significant expression of endothelial/vascular markers (**S6 Fig**) further confirming our observation (**Fig. 3**) about the role of late RA addition in limiting the induction of these pLPM derivatives. Moreover, the absence of *NKX2-1* and *FGF10* transcripts in our dataset (**S6 Fig**), together with the expression profile of these cells, argues against their identity as respiratory mesenchyme, which is also marked by co-expression of TBX4 and TBX5 [67].

**Fig 4.**
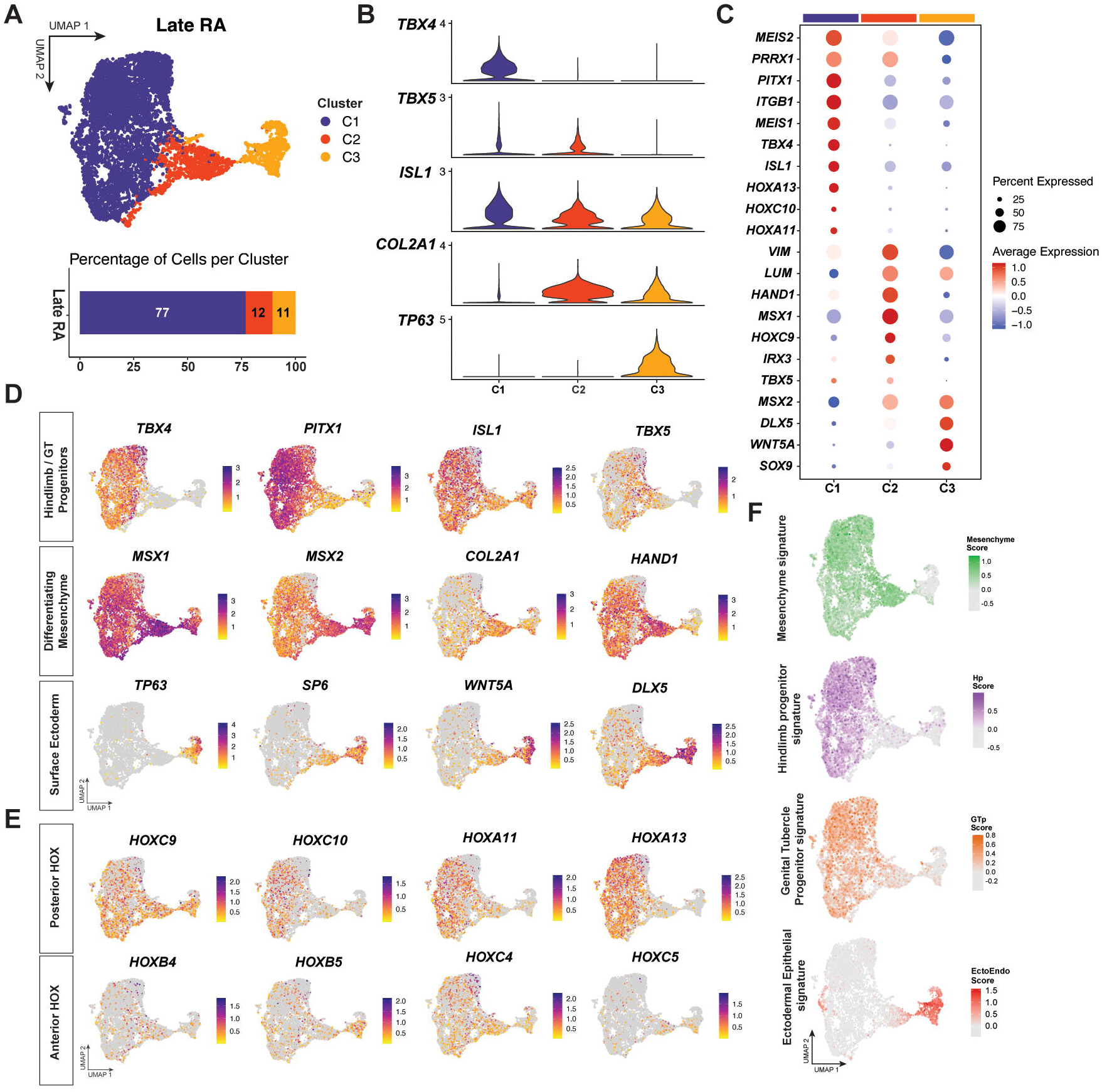
Single cell RNA sequencing reveals the co-differentiation of HGTp derivatives and ectodermal cells. **(A)** UMAP visualisation of sc-RNA seq data from late RA treated cells collected on day 10 of differentiation. Cells were divided into 3 clusters indicated by different colours. Below: Bar graph showing the percentage of total cells per cluster. **(B)** Stacked violin plots showing the expression (log-normalized) of key lineage markers enriched across identified cell clusters. **(C)** Dot plot showing the proportion of cells per cluster expressing the indicated marker genes. The colour bar indicates the average log-normalized expression values. **(D, E)** Feature UMAPs showing the expression of representative marker genes (log-normalized). **(F)** Projection of indicated marker signature scores on the UMAP. Cells coloured by the strength of their marker signature.

Cells in cluster 2 comprised a smaller population (12%) enriched for extracellular matrix and mesenchymal differentiation-associated transcripts, including *MSX1/2, COL2A1*, *LUM* and *VIM* (**Fig 4B-E**; **S8 Table**). The presence of these transcripts indicates further differentiation of cluster 1 progenitors toward a connective tissue/chondrogenic trajectory resulting from sustained BMP exposure, in line with previous reports on BMP treatment of hPSC-derived LPM cells [23,68]. The strong enrichment of *HAND1*, a BMP target [38,39], supports this interpretation. These cells nonetheless retained evidence of their origin: *ISL1* was maintained in 68% and *TBX5* in 39% of cells, whereas only 2% showed the *TBX4*⁺I*SL1*⁻ profile characteristic of a hindlimb trajectory, indicating that cluster 2 derives from the genital tubercle-biased fraction of cluster 1 rather than from hindlimb progenitors. *TBX4*, by contrast, was detected in only 10% of these cells (**S5 Fig**). As *TBX4* is maintained in genital tubercle mesenchyme *in vivo* [6,15,69], we interpret its loss here not as a feature of genital tubercle differentiation but as a departure from the appendage-field progenitor state as cells enter a general connective tissue programme .

Finally, cluster 3 (11% of total cells) displayed a strong surface ectoderm/epithelial cell signature as evidenced by the enrichment of transcripts such as *TP63, SP6, DLX5/6* and *WNT5A,* which are also expressed in both the limb and genital tubercle [67,70–80] (**Fig 4B-D** and **S7A Fig; S8 Table**). Immunofluorescence analysis revealed that these TP63^+^ epithelial cells surrounded an ISL1^+^ mesenchymal population (**S7B Fig**). This surface ectoderm/epithelial entity was already present at D5 of differentiation, comprising 42% of the total population (**S7C Fig**); late RA treatment reduced their abundance (18%), concomitant with an increase in the proportion of ISL1⁺ cells (85% from 74% at D5) (**S7C Fig**). No genital tubercle endoderm-associated markers were detected while the presence of apical ectodermal ridge-like *FGF8/TFAP2A/TP63/DLX* co-expressing cells was minimal (**S7D, S7E Fig**).

To further support the cluster annotations, gene signature scores were calculated for mesenchymal, ectodermal/epithelial, hindlimb and genital tubercle mesenchyme populations using established marker genes (**S9 Table**). Signature scores were projected onto the UMAP to assess their distribution across the identified cell populations (**Fig 4F**). Clusters 1 and 2 exhibited high mesenchymal signature scores, whereas cluster 3 was predominantly enriched for the ectodermal/epithelial signature, while also including a minor mesenchymal component (**Fig 4F**). Hindlimb and genital tubercle mesenchyme signatures predominantly marked cluster 1 and showed substantial overlap, reflecting the shared molecular features of these closely related progenitor populations (**Fig 4F**).

Collectively, these findings indicate that late activation of RA signalling largely promotes progression toward HGTps/genital tubercle mesenchymal derivatives that self-organise with an ectodermal/epithelial cell layer while limiting the emergence of pLPM-derived endothelial lineages.

### In vitro HGTp cultures resemble human embryonic pLPM-derived populations

To refine the annotation of our *in vitro*-derived cell populations, we mapped the scRNA-seq data onto the publicly available Human Embryogenesis Spatiotemporal Transcriptomic Atlas (HESTA) [81,82]. This resource includes spatial transcriptomic data from 77 sagittal sections of 13 human embryos spanning Carnegie Stages (CS) 12–23, capturing the development of key organs and cell populations relevant to our study. For label transfer analysis, four reference datasets representing the cloaca, limb bud, mesenchyme, and surface ectoderm were selected and integrated to generate an *in vivo* reference atlas (**Fig 5A**). A cardiac reference dataset was also included as a negative control to help distinguish between cardiac and HGTp *TBX5+* derivatives. Each *in vitro* cell was assigned to the reference population for which it received the highest prediction score, and the distribution of predicted identities was examined across the previously defined *in vitro* clusters. Overall, the predicted reference identities showed concordance of the posterior appendage/HGTp-associated cell types (cloaca, limb bud, mesenchyme) with the *in vitro* cluster annotations, supporting transcriptional correspondence between our hPSC-derived populations and their human *in vivo* counterparts (**Fig 5B**). Prediction scores were generally high, with particularly strong scores observed for the mesenchymal populations (**Fig 5C, Table S10**). In contrast, the hPSC-derived populations showed no strong transcriptional similarity to the heart-associated reference signature as only 0.04% of *in vitro* cells were assigned a cardiac identity (**Fig 5B, Table S10**).

**Fig 5.**
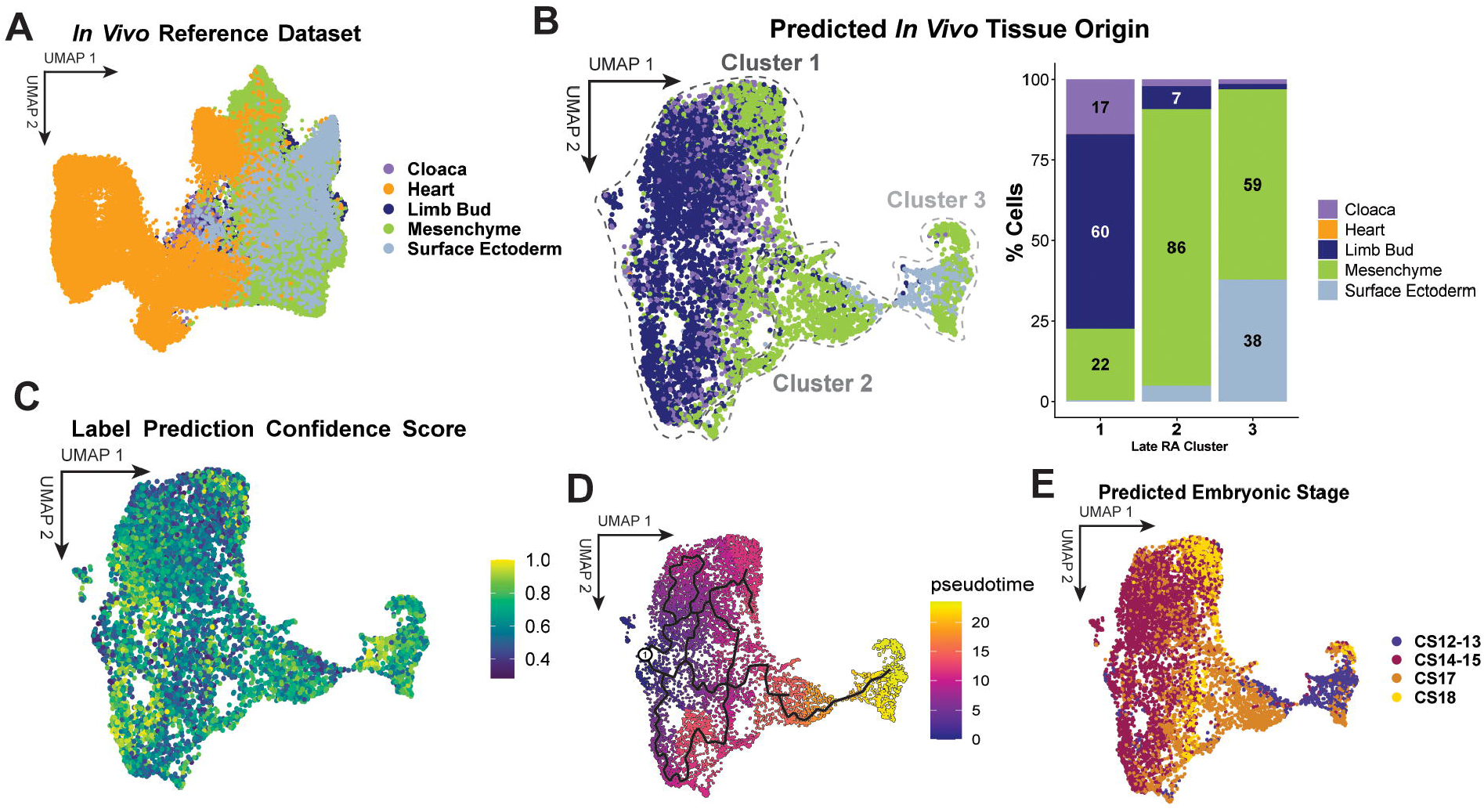
Comparison of in vitro-derived HGTps to their human embryonic counterparts. **(A)** UMAP plot of the integrated in vivo reference dataset coloured by embryonic tissue type (cloaca, limb bud, mesenchyme, heart and surface ectoderm). **(B)** UMAP visualisation showing the predicted in vivo tissue origin mapping for the in vitro dataset. **(C)** UMAP plot showing the label prediction confidence scores assigned to in vitro cells, gradient-coded from low (purple) to high (yellow) confidence. **(D)** Pseudotime trajectory reconstruction using Monocle3. The circled number (1) indicates the developmental trajectory root. **(E)** UMAP plot displaying the inferred human embryonic stage (Carnegie Stages CS12–CS18) of the in vitro-derived cells.

Pseudotime analysis using Monocle 3 [83] inferred a transcriptional trajectory extending from the mesenchymal-like progenitor population in cluster 1 towards the more committed mesenchymal populations represented by clusters 2 and 3, an ordering that is consistent with progressive differentiation (**Fig 5D**). To further contextualise these populations within human embryonic development, we compared their transcriptional profiles with HESTA-derived signatures across different Carnegie Stages. Cells within each *in vitro* cluster showed correspondence to multiple developmental stages rather than a single discrete time point, indicating heterogeneity in their developmental-state signatures (**Fig 5E**). The HGTp-enriched cluster 1 showed strongest correspondence with CS14–15 samples, consistent with a relatively early mesenchymal progenitor-like state, whereas cluster 2, enriched for more differentiated mesenchymal cells, showed the strongest correspondence with CS17 (**Fig 5E**).

Together, these analyses provide an independent human *in vivo* benchmark for our scRNA-seq dataset and indicate that our hPSC-derived cell populations exhibit transcriptional features corresponding to human embryonic appendage mesenchymal populations.

### Developmental potential of hPSC-derived HGTps in vivo

We next sought to determine the *in vivo* developmental potential of hPSC-derived HGTps. To this end, we generated HGTps/ genital tubercle progenitor cultures from SFCi55-ZsGr iPSCs, which are marked by constitutive expression of a ZsGREEN fluorescent reporter [60], using our 10-day “late RA” differentiation protocol (**Fig 3** and **4**). D10 cells were collected and cultured in suspension for a further 24 hours to generate spheroids of the appropriate size for grafting (**Fig 6A; S8A-B Fig**). These spheroids retained their molecular character as immunofluorescence analysis confirmed robust expression of the pLPM/HGTp/genital tubercle mesenchymal markers, HAND1, ISL1, and PITX1 and the presence of a minor TP63^+^ epithelial subpopulation (**S8C Fig**). Control spheroids of similar size, lacking expression of these markers, were also produced from undifferentiated hPSCs (**S8 Fig**).

**Fig 6.**
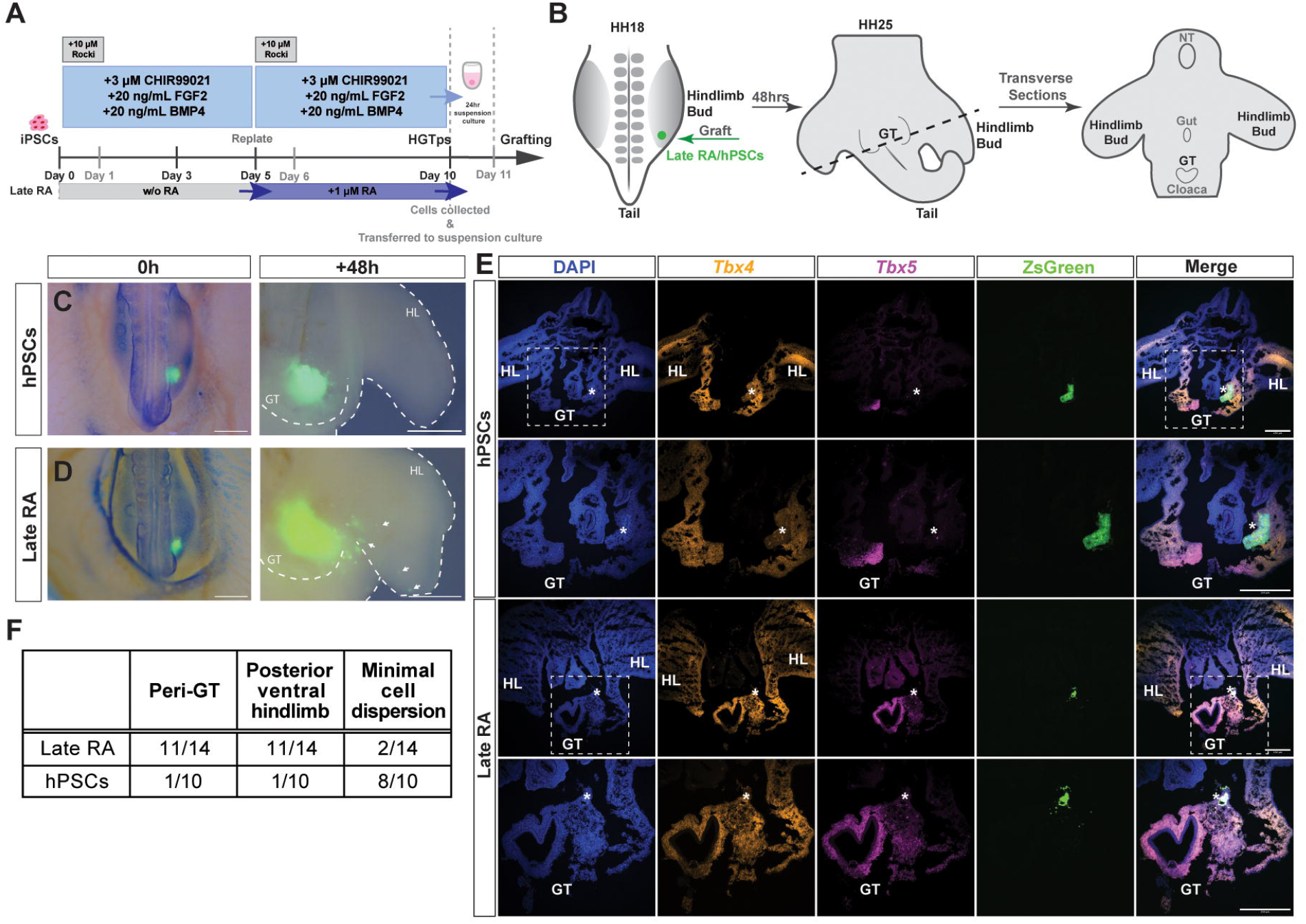
Grafting 3D human spheroids into chicken embryos. **(A)** Schematic of the treatment conditions used to generate spheroids from hPSC-derived HGTPs and hPSCs. **(B)** Experimental schematic depicting the site of spheroid grafting in HH18 chick embryos, 48-hour culture to stage HH25, and subsequent transverse sectioning through the genital tubercle (GT) and hindlimb bud regions. **(C)** Bright-field images of a representative chick embryo at 0 hours (left) and 48 hours (right) after grafting of a control ZsGREEN-positive hPSC spheroid. Scale bars represent 500 µm. **(D)** Bright-field images of a representative chick embryo at 0 hours (left) and 48 hours (right) after grafting of a ZsGREEN-positive hPSC-derived HGTP spheroid. Scale bars represent 500 µm. **(E)** Transverse sections of stage HH25 chick embryos analysed by HCR RNA-FISH, showing integration of donor ZsGREEN⁺ human cells (green) within host tissue expressing Tbx4 (orange) and Tbx5 (magenta). Magnified regions corresponding to the peri-genital tubercle (GT) area marked by a box are also shown. Cell nuclei are counterstained with DAPI (blue). Scale bars=250 µm. **(F)** Scoring of the grafts following whole mount embryo imaging, based on donor cell distribution within the peri-genital tubercle (GT) and posterior ventral hindlimb area.

Next, we grafted the hPSC-derived spheroids (HGTp and control) into a discrete territory within the pLPM located immediately posterior to the hindlimb bud and dorsal to the tailbud of Hamburger Hamilton 18 (HH18) chick embryos **(Fig 6B**). Previous fate mapping experiments revealed that this region harbours cells destined to contribute primarily to the genital tubercle area after the body wall closes at the ventral midline, but also to the posterior aspect of the hindlimb [8]. Human donor cell engraftment within host chick tissue was assessed 48 hours after transplantation. Following embryo sectioning, the identity and anatomical localisation of key structures, including genital tubercle mesenchyme, cloacal endoderm, and hindlimb tissue, were determined by HCR RNA-FISH using representative host markers (*TBX4, TBX5,* and *SHH*) **(Fig 6B; S9A Fig**). The majority (n=8/10) of engrafted control hPSC spheroids consistently failed to integrate into host tissue after 48 hours; instead, they formed condensed cell masses that were often extruded from the body at the nascent genital tubercle region (**Fig 6C, E, F**). By contrast, most (n=11/14) of the donor human ZsGREEN^+^ HGTp-derived spheroid grafts incorporated into the host peri-cloacal mesenchyme tissue within the genital tubercle-forming territory (**Fig 6C, E, F; S9B Fig**). The integration of human donor cells within this region was further confirmed by HCR, which revealed the presence of ZsGREEN⁺ cells within *TBX4/TBX5*-positive expression domains. (**Fig, 6E**). We also observed cells within the posterior part of the hindlimb bud that appeared migratory in nature, as indicated by their dispersed distribution (n=11/14) (arrowheads in **Fig 6D**). These findings demonstrate that *in vitro*-derived human HGTps exhibit positional competence when placed in a genital tubercle-fated region in the chick embryo.

## Discussion

Here, we describe a strategy that diverts hPSCs from an NMP trajectory to generate pLPM/HGTps that self-organise beneath an epithelial layer and show positional competence in xenografts.

### Generation of human pLPM/HGTps

Our BMP/WNT/FGF signalling regime generates a highly enriched pLPM/HGTp population, as indicated by induction of posterior HOXPG (*10–13*)⁺ lumbosacral genes and the emergence of *ISL1 TBX4* cells[1,6,12]. We show that high BMP activity promotes transcriptomic changes reminiscent of the trunk-to-tail transition and pLPM emergence. Although a role for Tgfb/BMP family ligands in mediating the Tgfbr1-dependent axial patterning of pLPM cells has been previously suggested in the mouse [3], our study presents, to our knowledge, the first direct evidence that BMP4 can functionally mimic the action of GDF11, a well-established regulator of the trunk-to-tail transition [1–3]. Gdf11, a member of the Tgfb/BMP superfamily, acts through Tgfbr1 to control posterior axial identity, and our findings suggest that BMP4 engages an overlapping signalling logic to promote pLPM specification. Although BMP4 and GDF11 preferentially activate SMAD1/5/8 and SMAD2/3, respectively, GDF11 has also been reported to activate SMAD1/5/8 in a cell context-dependent manner, while interactions between BMP- and TGFβ-associated SMAD pathways can integrate distinct upstream signals [84–86]. Consistent with this possibility, we observe induction of endogenous GDF11 following BMP4 treatment (**Fig. 2C**). Moreover, both Bmp4 and Gdf11 operate in cross-regulatory feedback loops involving Isl1 to specify pLPM-derived lineages in the mouse, including hindlimb and urogenital tissues [1,2,12,16]. Additional work in chick embryos demonstrates that an increasing medial-to-lateral BMP4 gradient specifies distinct LPM fates [87,88], with the genital tubercle arising at the most-lateral position [7,8,11]. Together, our findings reinforce the idea that high BMP signalling drives trunk-to-tail transition-like transcriptome changes and pLPM/HGTp fate acquisition and reveal that these processes are conserved in human development.

### Acquisition of genital tubercle progenitor identity

We show that the induction of *TBX5* accompanies expression of posterior *HOXA–D* PG (*10– 13*) genes, key regulators of early genital tubercle development [5]. This relationship indicates that *TBX5*⁺ cells do not represent aLPM-derived lineages such as forelimb or cardiac mesenchymal progenitors. Accordingly, *Tbx5* expression initiates in the mouse genital tubercle mesenchyme later than *Tbx4* [51], closely mirroring our bulk RNA-seq data. Interestingly, recent scRNA-seq data identifies *TBX4^+^TBX5^+^* cells in the human genital tubercle at a comparable early stage (Carnegie Stage 13) [15]. Our protocol also generates *TBX4 TBX5* cells, indicating that this defining feature of human genital tubercle specification is reproduced in our cultures.

Our scRNA-seq data infers differentiation trajectories from pLPM/HGTps and indicates that mesenchymal cells progressively acquire genital tubercle rather than hindlimb character. Cluster 1, comprising HGTps together with cells acquiring a genital tubercle character, contains a subset of *TBX5*⁺ cells (+/- for *ISL1* and/or *TBX4*), consistent with early commitment toward a genital tubercle progenitor state. By contrast, Cluster 2 is enriched in markers of mesenchymal differentiation including collagens and predominantly contains *TBX5*⁺ and/or *ISL1*⁺ cells suggesting the adoption of a general connective tissue differentiation route, while retaining markers of their genital tubercle progenitor identity. This is likely to be driven by BMP signalling in line with previous reports examining the effect of BMP treatment on hPSC-derived LPM cells [23,68]. Therefore, manipulating BMP signalling at later stages of the protocol is an obvious route to improving regional identity. Importantly, comparison of our scRNA-seq data with spatial transcriptomic data from human embryos [81,82] indicated that the hPSC-derived HGTp populations and their mesenchymal derivatives resemble their corresponding *in vivo* counterparts at the transcriptional level. Importantly, only 0.04% of cells were assigned a cardiac identity, despite *TBX5* being prominently expressed in the cardiac lineage as well as in the genital tubercle. This low proportion indicates that the mapping is not biased by *TBX5* expression alone. Nonetheless, we note that marker-based inference of cell identity across species, and between *in vitro* and *in vivo* systems, has inherent limitations. Although we have anchored our annotations to human reference datasets [15,81,82], some markers were originally characterised in mouse, and differences in expression between species or between culture and embryo may remain. In addition, our HGTps correspond to a stage at which the human genital tubercle has not yet subdivided into its derivative populations, so no stage-matched *in vivo* population is available against which cluster 2 can be benchmarked directly.

Our results indicate that the late stimulation of RA signalling refines the differentiation of HGTps toward a genital tubercle mesenchymal fate at the expense of pLPM endothelial/vascular derivatives. Although the Cyp26a1-dependent suppression of posterior RA activity is required for trunk and lumbosacral tissue formation[1,4], the RA synthesising gene, *Aldh1a2*, is expressed later in mouse genital tubercle cells[46,64]. This temporal shift is consistent with a critical role for RA in specifying pLPM/HGTp derivatives, likely through mesenchymal–epithelial interactions [64]. By contrast, early RA stimulation at the exit from pluripotency, blocked the trunk-to-tail-like transition toward pLPM.

### Self-organisation of HGTps and their mesenchymal derivatives with an ectodermal epithelium

Our scRNA-seq analysis identified a distinct subpopulation with ectodermal/epithelial characteristics that co-emerges with pLPM cells, consistent with studies showing that BMP signalling directs hPSCs toward non-neural ectodermal fates [89,90]. This suggests that differential interpretation of BMP activity drives the concurrent emergence of distinct mesenchymal and surface ectoderm/epithelial lineages, which subsequently self-organise into spatially distinct domains. Reciprocal interactions between mesenchymal and epithelial compartments are essential for hindlimb and external genitalia formation *in vivo*. Accordingly, defining the spatial organisation and functional interplay of these cell types in our system will provide a powerful platform for modelling human posterior appendage development and probing the origins of congenital or environmentally induced malformations. However, determining how these populations assemble into embryonic-like architecture will also require bioengineered 3D organoid platforms, tissue explants and high-multiplex spatial transcriptomics [91,92].

### Human HGTps contribute to the genital tubercle forming region of the chick embryo

Our results demonstrate that spheroids generated from D10 hPSC-derived HGTps contribute to the external genital-forming territory when xenografted into pLPM between the nascent hindlimb and genital tubercle forming regions of HH18 chick embryos. These findings reflect the observation that these spheroids maintain their HGTp/genital tubercle identity (HAND1, ISL1, PITX1), consistent with molecular and positional parity with the host tissue. By contrast, control hPSC-derived spheroids, which do not express these markers, are extruded from the tissue as condensed cell masses.

HGTp-derived spheroids also contributed cells to the posterior region of the hindlimb. However, their dispersed distribution suggests they represent a distinct migratory subpopulation rather than hindlimb progenitors. These findings suggest that our differentiation protocol produces genital tubercle rather than hindlimb mesenchyme-biased progenitors in line with our scRNA-sequencing data.

Taken together, our work provides a human system in which the divergence of the hindlimb and genital tubercle lineages can be followed from pluripotency. Comparison with human embryo datasets places these HGTps within the appropriate pLPM lineage at Carnegie stages 14–17, and their behaviour on xenografting indicates positional competence in an embryonic environment. Future work could address when and how fate becomes fixed between the hindlimb and genital tubercle lineages, which the spectrum of states we observe suggests is gradual rather than switch-like. A second question is whether regional identity can be better controlled, given the general connective tissue route that sustained BMP exposure appears to impose. Resolving both will be necessary for modelling the congenital conditions that affect the hindlimbs and external genitalia together, and for any tissue engineering application that depends on reproducible, fate-committed genital tubercle progenitors.

## Methods

### hPSC culture and differentiation

The human induced pluripotent stem cell line 113-CR-3 (male) was employed in all experiments^30^. Cells were karyotyped and tested regularly for mycoplasma using Mycostrip Mycoplasma detection kit (InvivoGen, rep-mys-50) and expression of pluripotency markers. Copy Number Variation (CNV) analysis was carried out by Stem Genomics using the iCS-digital PSC method, which revealed the presence of a chromosome 20q gain in the 113CR3 cell line, a common aberration found in almost 20% of hPSCs tested worldwide [93]. Experiments were repeated using a second, karyotypically normal iPSC line, SFCi55-ZsGr iPSCs [60] as well as the embryonic stem cell line WA09/H9 [61]. The use of hESCs in this project has been approved by the Human Embryonic Stem Cell UK Steering Committee (SCSC15-23). Undifferentiated cells were cultured routinely in feeder-free conditions in mTeSR1 (Stem Cell Technologies, 85850) medium on Geltrex LDEV-Free reduced growth factor basement membrane matrix (Thermo Fisher Scientific, A1413202)-coated plates. Cells were passaged every 4-5 days when the confluency reached approximately 80% using ReLeSR (Stem Cell Technologies, 100-0484).

Differentiation experiments were conducted on vitronectin (VTN-N) (Thermo Fisher Scientific, A31804)-coated culture plates. Cells were cultured in N2B27 basal medium containing 50:50 Dulbecco’s Modified Eagle’s Medium (DMEM) F12 (Sigma-Aldrich, D6421)/Neurobasal medium (Gibco, 21103049) and 1xN2 supplement (Gibco, 17502001), 1xB27 (Gibco, 17504001), 1xGlutaMAX (Gibco, 35050061), 1xMinimum Essential Medium Non-Essential Amino Acids (MEM NEAA) (Gibco, 11140050), 2-mercaptoethanol (50 μM, Gibco, 31350010). NMP differentiation was conducted as previously explained[34]. For LPM differentiation, hiPSCs after reaching 70-80% confluency, were dissociated using TrypLE (Gibco, 12563029) and seeded at a 60,000 cells/cm^2^ density. The Rho-associated coil kinase (ROCK) inhibitor Y-27632 2HCl (Adooq Biosciences, 10 μM, A11001) was added for the first day of differentiation to support survival of the cells plated as a single-cell suspension. The N2B27 medium was supplemented with CHIR99021 (3 μM, Tocris, 4423), FGF2 (20 ng/ml, R&D Systems, 233-FB-500/CF), BMP4 (20 ng/mL, Thermo Scientific, PHC9531) and either with RA (1 μM, Sigma-Aldrich, R2625) or without RA for control. Cultures were fed with fresh supplemented media on the second day, and cells were cultured for 3 days. For testing the effect of early RA treatment, the culture was extended to 5 days and cells were fed with N2B27 supplemented with CHIR99021 (3 μM), FGF2 (20 ng/ml), BMP4 (20 ng/mL) and RA (1 μM) in every other day after the first day. For testing the effect of late RA treatment, RA (1 μM) was excluded from the medium. ROCK inhibitor Y-27632 2HCl (10 μM) was added for the first day only. For longer term differentiation, cells were cultured in N2B27 supplemented with CHIR99021 (3 μM), FGF2 (20 ng/ml), BMP4 (20 ng/mL) and treated either with RA (1 μM) (for early RA treatment) or left untreated (for late RA condition). ROCK inhibitor Y-27632 2HCl (10 μM) was added for the first day. 24h later, media was replaced removing the ROCK inhibitor and cultured for 5 days. On day 5, cells were dissociated using TrypLE and re-plated at a 80,000 cells/cm^2^ density in N2B27 supplemented with CHIR99021 (3 μM), FGF2 (20 ng/ml), BMP4 (20 ng/mL) and RA (1 μM) for both late RA and early RA conditions or left untreated for control group. ROCK inhibitor Y-27632 2HCl (10 μM) was added on day 5 during replating. Cells were fed in every other day after the day 1 and day 6.

### Spheroid preparation and culture

Day 10 late RA-treated cells and hPSCs, used as controls, were dissociated into single cells using TrypLE. Cells were seeded into 96-well low-attachment plates at a density of 5,000-6,000 cells per spheroid. For the RA-treated group, cells were cultured in N2B27 medium supplemented with CHIR99021 (3 μM), FGF2 (20 ng/mL), BMP4 (20 ng/mL), RA (1 μM), and Y-27632 2HCl (10 μM). Control hPSCs were seeded in mTeSR medium containing Y-27632 2HCl (10 μM). Plates were centrifuged immediately after seeding to promote 3D spheroid formation. Cells were incubated for 24 hours prior to grafting.

### Quantitative real-time PCR

Total RNA was extracted from the cells using the total RNA purification kit (Norgen Biotek, 17200) according to the manufacturer’s instructions. RNA concentration and purity were assessed using a NanoDrop Lite spectrophotometer (Thermo Fisher Scientific). The cDNA synthesis was performed using the High-Capacity cDNA Reverse Transcription kit (Thermo Fisher Scientific, 4368814). The quantitative real-time PCR was carried out using the QuantStudio 12K Flex Real-Time PCR System (Applied Biosystems) using PowerUp SYBR Master Mix (Thermo Fisher Scientific, A25780). The relative expression levels were calculated with the comparative threshold cycle (ΔΔCT) method and normalised to either GAPDH or TBP expression. Primers are listed in Supplementary Table 11. Graphs were generated with GraphPad Prism (GraphPad Software), which was also used for statistical analysis.

### Immunocytochemistry and image analysis

Cells were fixed in 4% paraformaldehyde (VWR, J61899.AP) for 10 min at room temperature (RT), then rinsed three times with PBS. The cells then permeabilised and blocked with blocking buffer containing 0.1% Triton X-100 (Sigma-Aldrich, X100-500 ML) and 1% bovine serum albumin (Sigma-Aldrich, A7906-100G) in PBS for 1 hour at RT. Subsequently, primary antibodies were diluted in blocking buffer, and cells were incubated with primary antibodies overnight at 4 °C and then washed with PBS 3 x 5 min. Following primary antibodies were used; TBXT (Abcam, ab209665, 1:1000), ISL1 (DSHB, 39.4D5, 1:200), HAND1 (R&D Systems, AF3168, 1:100), PITX1 (Novus Bio, NBP1-88644, 1:500), HOXC9 (Abcam, ab50839, 1:200), KDR (R&D Systems, AF357, 1:1000), TP63 (Abcam, ab124762, 1:200). The secondary antibodies conjugated to Alexa fluorophores (Invitrogen) diluted in blocking buffer and the cells were incubated with secondary antibodies for 2 h in the dark at RT and washed 3 x 5 min with PBS. Secondary antibodies employed: Donkey anti-Rabbit IgG (H+L) Highly Cross-Adsorbed Secondary Antibody, Alexa Fluor488 (Invitrogen, A-21206, 1:500), Donkey anti-Mouse IgG (H+L) Highly Cross-Adsorbed Secondary Antibody, Alexa Fluor 594 (Invitrogen, A-21203, 1:500), Donkey anti-Goat IgG (H+L) Cross-Adsorbed Secondary Antibody, Alexa Fluor 647 (Invitrogen, A-21447, 1:500), Donkey anti-Rat IgG (H+L) Cross-Adsorbed Secondary Antibody, DyLight 650 (Invitrogen, SA5-10029, 1:500), Donkey anti-Rabbit IgG (H+L) Highly Cross-Adsorbed Secondary Antibody, Alexa Fluor 594 (Invitrogen, A-21207, 1:500), Donkey anti-Mouse IgG (H+L) Highly Cross-Adsorbed Secondary Antibody, Alexa Fluor 488 (Invitrogen, A-21202, 1:500), Donkey anti-Goat IgG (H+L) Cross-Adsorbed Secondary Antibody, Alexa Fluor 594 (Invitrogen, A-11058, 1:500). Cell nuclei were stained with DAPI (Thermo Fisher Scientific, 62248) diluted in PBS 1:1200. Immunofluorescent microscopy was conducted with InCell Analyser 2200 system (GE Healthcare) or confocal microscopy with W1 Spinning Disc Confocal (Nikon). Images were processed in Fiji[94] and image quantifications were performed using CellProfiler[95] with identical brightness/contrast settings to allow comparison between different conditions. The negative threshold was set using a sample incubated with secondary antibody only. Area and roundness values of the hPSC-derived 3D models were measured using Fiji [94], Measure function.

### Whole mount immunostaining

Spheroids were collected and fixed in 4% paraformaldehyde for 30 min at RT, followed by three washes with PBS. Samples were permeabilised in PBS containing 0.3% Triton X-100 for 30 min at RT and then blocked in PBS containing 1% bovine serum albumin and 0.1% Tween-20 (Sigma-Aldrich, P9416) for 2 hours at RT. Primary antibodies (as listed above) were diluted in blocking buffer and spheroids were incubated overnight at 4 °C. Samples were washed with PBS (3 × 5 min) and incubated with Alexa fluorophore–conjugated secondary antibodies (as listed above) diluted in blocking buffer for 2 hours at RT protected from light. Following washes with PBS (3 × 5 min), nuclei were stained with DAPI (1 µg/mL in PBS) for 10 min at RT in the dark. Spheroids were washed again with PBS (3 × 5 min) and imaged using confocal microscopy with a W1 Spinning Disc Confocal system.

### Bulk RNA-sequencing

#### RNA Isolation, library preparation and bulk RNA-seq

Total RNA was separately isolated from three replicates of the differentiated cells from each condition at the indicated time points and hiPSCs as described above using the total RNA purification kit (Norgen Biotek, 17200) according to the manufacturer’s instructions. RNA concentration and purity of each RNA sample was evaluated using the BS RNA kit (Invitrogen, Q10210) and the Qubit 3.0 Fluorometer (Life Technologies, Q33216). Samples were then submitted to Novogene for library preparation and bulk-RNA seq. Library preparation and sequencing were performed after sample quality control by Novogene using an Illumina NovaSeq X Plus Series, PE150 platform. In-house perl scripts were used to process the raw data of FASTQ format and clean data were calculated by removing reads containing adapter, reads containing poly-N and low-quality reads from raw data. Analysis was completed by Novogene with reference genome (Homo Sapiens [GRCh38/hg38]).

#### Read mapping, quantification of gene expression level, DEGs and GO enrichment analysis of bulk RNA-seq data

Mapping to the reference genome was done by Hisat2 (v2.0.5) mapping tool[96]. Quantification of gene expression level was done using featureCounts (v1.5.0-p3)[97] and Fragments Per Kilobase of transcript per Million mapped reads (FPKM) of each gene was calculated based on the length of the gene and read counts mapped to that gene. FPKM values used for normalisation, and Principal Component Analysis. Differential expression analysis was performed with DESeq2 (v1.20.0)[98]. Genes with an adjusted p-value<=0.05 found by DESeq2 were assigned as differentially expressed. Functional analysis was performed using clusterProfiler (v3.8.1)[99] R package for enrichment analysis, including GO Enrichment, KEGG and Reactome database Enrichment. Statistical significance was evaluated using a hypergeometric test, with multiple testing correction applied using the Benjamini-Hochberg false discovery rate (FDR). Enriched terms with adjusted p-values < 0.05 were considered statistically significant. Plotting was done using R or NovoMagic.

### Single-cell RNA sequencing

#### Single cell preparation for sc-RNA seq

Cells were harvested on day 10 of differentiation and dissociated into a single-cell suspension using TrypLE, followed by centrifugation at 300 rcf for 5 minutes. The cell pellet was resuspended in STEM-CELL BANKER (AMSBIO) and filtered through a cell strainer to remove cell aggregates. Cell viability was assessed by Trypan Blue exclusion, and live cells were cryopreserved at a density of 1×10^6^ cells/mL in STEM-CELL BANKER until further processing.

#### Library generation and sequencing

Library preparation and single-cell RNA sequencing was performed by Active Motif using 10X Genomics Single Cell 3’ (v3) kit. Libraries were sequenced on Illumina NovaSeq 6000 platform as paired-end 91 bp reads. Approximately 10,000 cells were analysed at 50,000 reads/cells. Raw sequencing data were processed using the Cell Ranger pipeline (10x Genomics). Reads were aligned to the human reference genome GRCh38, and gene-cell count matrices were generated using default parameters.

#### Data processing and analysis

Filtered count matrices were imported into Seurat (v5.3.1)[100] for downstream analysis. Quality control measures were applied by excluding features fewer than 1000, greater than 10,000; counts fewer than 1000, greater than 50,000 and cells with over 15% mitochondrial gene expression (**S4 Fig**).

Standard pre-processing was performed using Seurat. Data were normalized using Seurat’s *NormalizeData* function with a scale factor of 10,000. Highly variable genes were identified using *FindVariableFeatures,* selecting the top 2,000 variable genes. Doublets were identified and removed using DoubletFinder (v2.0)[101], with an estimated doublet number of 478 cells. Subsequent analyses were performed on the remaining 7,814 singlet cells.

Following doublet removal, data were re-normalized and scaled using the same parameters, and principal component analysis (PCA) was performed using *RunPCA*. The first 12 principal components were used to construct a shared nearest-neighbor graph (*FindNeighbors*) and for clustering (*FindClusters*) at a resolution of 0.03. Cluster visualization was performed using Uniform Manifold Approximation and Projection (UMAP). Cluster-specific marker genes were identified using *FindAllMarkers.* Marker gene expression was visualized using the scCustomize package (v3.2.2)[102] with *FeaturePlot_scCustom*, displaying positive expression values on a continuous colour scale. Gene signature scores were calculated using manually curated marker gene sets derived from published studies of mesenchymal, surface ectoderm/epithelial, hindlimb progenitor, and genital tubercle progenitor (Table S9) using *AddModuleScore* function. Pseudotime analysis applied using Monocle (v3) [103].

#### Label transfer and cross dataset annotation

Publicly available Human Embryogenesis Spatiotemporal Transcriptomic Atlas (HESTA) [81,82] scRNA-seq data from human embryos was mapped onto the *in vitro*-generated data using Seurat’s label transfer workflow. Five reference datasets from different stages were used: limb bud, cloaca, mesenchyme, surface ectoderm and heart. Reference datasets were merged using Seurat. Top 3,000 highly variable genes were identified using *FindVariableFeatures,* vst method (v2.0) [100]. Data were processed using *ScaleData* and PCA analysis was performed using *RunPCA*. Harmony (v2.0.5) [104]. was used for integration and batch correction. Label transfer was applied using FindTransferAnchors and TransferData functions of Seurat. Cell mappings were visualised on UMAPs using DimPlot.

### Chick husbandry and spheroid grafting

Wild-type (Medeggs, Norfolk, UK) Bovan Brown eggs were incubated and the embryos staged according to Hamburger Hamilton stages. For spheroid grafting a sharpened tungsten needle was used to cut a slit on the posterior 1/5 of the lateral plate mesoderm at the hindlimb level. A GFP spheroid, either late-RA or hPSC, was then grafted into the opening. The grafted embryos were covered and left to develop for 48h before collection for RNA-fluorescence in situ hybridisation. Grafts were imaged at 0h and immediately after collection to check position of the grafts using a LeicaMZ16F microscope and Flexacam C3 Camera with LAS X 3.10.2.30243 imaging software.

### RNA fluorescence in situ hybridization with amplification by Molecular Instruments Hybridization Chain Reaction v3.0

Grafts were fixed overnight in 4% PFA at 4°C, then washed in PBS and dehydrated in a methanol-PBT series. Dehydrated grafts were stored in methanol at -20°C before RNA fluorescence in situ hybridization by Molecular Instruments Hybridization Chain Reaction (HCR) v3.0 [105] was performed. For HCR the samples were rehydrated in a methanol-PBT series, treated proteinase K for 25 minutes, followed by post-fixing in 4% PFA for 20 minutes. The samples were then bleached to remove autofluorescence in a 3% H_2_O_2_/20 mM NaOH/PBS solution for 30 minutes on ice. The samples were further washed in PBT and 5X SSCT. Samples were pre-hybridized in HCR Probe Hybridization Buffer (v3.0) for at least 1hour at 37 °C before addition of probe solution overnight at 37 °C. All probes were acquired from Molecular Instruments Inc. and part of their Infinite Catalog. The probe solution was prepared by adding 1 μl of 1μM probe per 100 μl of HCR Probe Hybridization Buffer (v3.0). The following day the samples were washed in HCR Probe Wash Buffer (v3.0) and 5X SSCT before pre-amplifying in HCR Amplification Buffer (v3.0) for at least 1 hour at room temperature. Samples were then incubated in fluorescently labelled amplifiers in HCR Amplification Buffer overnight at room temperature in the dark. The amplifiers were prepared by heat-shocking 2μl of each amplifier pair at 95 °C for 90s and cooling for 30 minutes at room temperature before adding to 100 μl of HCR Amplification Buffer (v3.0). Samples were then washed in 5X SSCT containing DAPI (1mg/ml, 1:1000). Samples were further fixed, incubated in 30% Sucrose (Sigma-Aldrich, S0389-500G) overnight and mounted in OCT (Fisher Scientific, 12678646) for cryosectioning (30 µm sections). HCRs were imaged on a Nikon W1 Spinning Disk Confocal Microscope (10X and 20X objective) with Nikon Elements Software.

## Data availability

The bulk RNA-sequencing data and single-cell RNA sequencing data generated in this study have been deposited in the Gene Expression Omnibus (GEO) under accession codes GSE318031, and GSE318032 respectively.

## Acknowledgements

This work was supported by the Medical Research Council (MRC) (MR/Y013476/1; AT) and the Biotechnology and Biological Sciences Research Council (BB/P000444/1; AT and X/014063-23-22; SU)(BB/Y012615/1; APP58087/UKRI2946; MT). We would like to thank Darren Robinson and Nick Van Hateren for help with imaging, which was performed at the Wolfson Light Microscopy Facility. We are also grateful to Luz Garcia-Alonso, Valentina Lorenzi, Roser Vento, Gi Fay Mok and Mark Dunning for help with sc-RNAseq analysis and Moises Mallo, Marysia Placzek, Filip Wymeersch, Minoru Takasato and Val Wilson for valuable discussions and critical reading of the manuscript. For the purpose of open access, the authors have applied a Creative Commons Attribution (CC BY) licence to any Author Accepted Manuscript version arising from this submission.

## Author contributions

**Conceptualisation:** SU, MT, AT

**Methodology:** SU, SSP, KC

**Formal analysis:** SU, SFP, MT, AT

**Investigation:** SU, SSP, KC

**Data curation:** SU, SSP

**Writing - Original Draft:** SU, SSP, MT, AT

**Writing - Review & Editing:** SU, SSP, MT, AT

**Supervision:** MT, AT

**Project administration:** MT, AT

**Funding acquisition:** MT, AT

## Supporting Information

**S1 Fig. Testing the effect of BMP stimulation during hPSC differentiation toward NMPs.**

**(A)** Schematic showing the key stages and markers associated with the transition of pLPM cells toward genital tubercle (GTps) and hindlimb mesenchyme progenitor (Hps) cells via HGTps [15,106]. Note that other pLPM derivatives are not included. **(B)** Top: Schematic of the treatment conditions used to test different BMP4 concentrations (10, 15, 20 ng/mL). Bottom: corresponding RT-qPCR expression analysis of key NMP and pLPM markers after 3 days of differentiation is shown below (*n* = 3 biologically independent experiments). Relative expression levels to iPSCs normalised to GAPDH gene expression (* p<0.05, ** p<0.01, *** p<0.001, mean ±SD) (One-way ANOVA and Tukey’s test). **(C)** Volcano plot of differentially expressed genes between BMP treated pLPM/HGTps vs untreated day 3 NMP cultures pLPM/HGTps on day 3.

**S2 Fig. Induction of HGT/genital tubercle markers following differentiation of SFCi55-ZsGr and H9 hPSCs.**

**(A)** RT-qPCR-based expression analysis of key pLPM/HGTp/genital tubercle markers after 5 days of SFCi55-ZsGr iPSC (top) and H9 hESC (bottom) differentiation (*n* = 3 independent experiments). Relative expression levels to iPSCs were normalised to TBP gene expression (* p<0.05, ** p<0.01, *** p<0.001, mean ±SD) (paired, two-tailed *t*-test). **(B)** Immunofluorescence analysis of the expression of HAND1 and PITX1 on day 5 of differentiation. Cell nuclei are counterstained with DAPI (blue). Scale bars represent 100 µm. Image analysis of the percentage of nuclei positive for PITX1 and HAND1 protein expression is also shown. Graph shows mean values (*n* = 2 independent experiments).

**S3 Fig. Examining the effect of distinct RA treatment regimes on differentiating SFCi55-ZsGr and H9 hPSCs.**

**(A)** RT-qPCR-based expression analysis of key HGTp/genital tubercle markers in day 10 SFCi55-ZsGr and **(B)** H9 untreated, late RA and early RA treated cells (*n* = 3 independent experiments). Relative expression levels to iPSCs were normalised to TBP gene expression (* p<0.05, ** p<0.01, *** p<0.001, mean ±SD) (One-way ANOVA and Tukey’s test).

**S4 Fig. Quality control of scRNA-seq analysis.**

**(A)** Data summary (unfiltered) and thresholds used for subsetting (dashed lines) the data during quality control. **(B)** Cell counts before and after quality control and filtering. **(C)** UMAP showing the doublets (478 cells) and singlets (7814 cells) detected by *DoubletFinder*.

**S5 Fig. HGTp/genital tubercle mesenchyme marker expression**

**(A)** UMAP showing the proportions of *ISL+/TBX5+/TBX4+* cells across the identified cell clusters. Below: Bar graph showing the percentage of cells per cluster calculated for each cluster. **(B)** UMAP visualisation of genital tubercle mesenchyme marker expression (*TBX4+TBX5+ISL1+*). **(C)** UMAP showing the proportions of *HOXB13+/TBX5+* cells across the identified cell clusters. Below: Bar graph showing the percentage of cells per cluster calculated for each cluster. **(D)** UMAPs showing the *TBX5+HOXPG4/5+* cells across identified cell clusters. Below: Bar graph showing the percentage of cells per cluster calculated for each cluster.

**S6 Fig. Expression of non HGT/GT markers**

Feature UMAPs showing the percentage expression of representative cardiac, endothelial and respiratory mesenchyme marker genes. Und. Undetected.

**S7 Fig. Ectoderm marker expression**

**(A)** Dot plot showing the percentage expression of key surface ectoderm markers for each cell cluster. The colour bar indicates the average log-normalized expression values. **(B)** Immunofluorescence analysis of the expression of TP63 and ISL1 on day 10 HGTps. Sequential z-stacks are shown (z=1 bottom, z=20 top). Scale bars represent 100 µm. **(C)** Immunofluorescence analysis of the expression of TP63 and ISL1 in day 5, and day 10 cells. Scale bars represent 100 µm. The percentages of nuclei positive for ISL1 and TP63 protein expression following image analysis are also shown on. Cell nuclei are counterstained with DAPI (blue). The graph shows mean values (n = 3 independent experiments) (* p<0.05, ** p<0.01, *** p<0.001, mean ±SD) (one way ANOVA and Tukey’s multiple comparison test). **(D)** Feature UMAPs showing the percentage expression of representative endoderm associated markers **(E)** UMAPs showing the proportions of apical ectodermal ridge markers.

**S8 Fig. Generation of HGTp spheroids from hPSCs**

**(A)** Bright-field images of spheroid models after 24 hours of incubation, prior to grafting. **(B)** Morphometric analysis of late RA-treated and hPSC spheroids including area (mm^2^) and roundness (n = 6 independent experiments) (* p<0.05, ** p<0.01, *** p<0.001, mean ±SD) (Welch’s two-sample t-test). **(C)** Immunofluorescence analysis of indicated HGTp/ectoderm markers in late RA-treated and hPSC spheroids. Cell nuclei are counterstained with DAPI (blue). Scale bars represent 100 µm.

**S9 Fig. Assessing the transplantation of hPSC-derived HGTps into chick embryos.**

**(A)** Wholemount (top row) and serial transverse sections (A–C, rows 2–4) of host chick embryos showing expression of *Tbx5* (magenta), *Tbx4* (orange), and *Shh* (green) detected by HCR RNA-FISH. Cell nuclei are counterstained with DAPI (blue). White lines on the wholemount image denote the planes of transverse sections A, B, and C. Scale bars= 250 µm. **(B)** Representative transverse sections of stage HH25 chick embryos analysed by HCR RNA-FISH, showing integration of donor ZsGREEN⁺ human cells (green) within host tissue expressing Tbx4 (orange) and Tbx5 (magenta). Cell nuclei are counterstained with DAPI (blue). Magnified regions corresponding to the peri-genital tubercle (GT) area marked by a box are also shown. White arrows show examples of integrated human donor cells. Scale bars= 250 µm.

**S1 Table. DEGs between day 3 BMP treated cells versus day 3 untreated cells**

**S2 Table. DEGs between day 5 cells versus day 3 cells**

**S3 Table. DEGs between day 5 cells versus iPSCs**

**S4 Table. GO Enriched terms on upregulated genes of day 5 BMP treated cells vs iPSCs**

**S5 Table. DEGs between day 10 late RA treated cells versus day 10 early RA treated cells**

**S6 Table. DEGs between day 10 late RA treated cells versus day 10 untreated cells S7 Table. DEGs between day 10 late RA treated cells versus day 5 cells**

**S8 Table. scRNA-seq cluster marker genes**

**S9 Table. Marker gene lists used to calculate signature scores.**

**S10 Table. Late RA best vs second best predictions**

**S11 Table. Primer Sequences used for quantitative real-time PCR RawData Table. Raw data used for generating the graphs shown in figures 1-3**.

## References

1. Jurberg AD, Aires R, Varela-Lasheras I, Nóvoa A, Mallo M. Switching Axial Progenitors from Producing Trunk to Tail Tissues in Vertebrate Embryos. Developmental Cell. 2013;25: 451–462. doi:10.1016/j.devcel.2013.05.009

2. Lozovska A, Korovesi AG, Dias A, Lopes A, Fowler DA, Martins GG, et al. Tgfbr1 controls developmental plasticity between the hindlimb and external genitalia by remodeling their regulatory landscape. Nat Commun. 2024;15: 2509. doi:10.1038/s41467-024-46870-z

3. Lozovska A, Casaca A, Novoa A, Kuo Y-Y, Jurberg AD, Martins GG, et al. Tgfbr1 regulates lateral plate mesoderm and endoderm reorganization during the trunk to tail transition. eLife. 2025;13: RP94290. doi:10.7554/eLife.94290.3

4. Wymeersch FJ, Wilson V, Tsakiridis A. Understanding axial progenitor biology in vivo and in vitro. Development. 2021;148: dev180612. doi:10.1242/dev.180612

5. Hintermann A, Bolt CC, Hawkins MB, Valentin G, Lopez-Delisle L, Ryan MM, et al. Co-option of an ancestral cloacal regulatory landscape during digit evolution. Nature. 2025;648: 109–116. doi:10.1038/s41586-025-09548-0

6. Naiche LA, Arora R, Kania A, Lewandoski M, Papaioannou VE. Identity and fate of *Tbx4* expressing cells reveal developmental cell fate decisions in the allantois, limb, and external genitalia. Developmental Dynamics. 2011;240: 2290–2300. doi:10.1002/dvdy.22731

7. Tschopp P, Sherratt E, Sanger TJ, Groner AC, Aspiras AC, Hu JK, et al. A relative shift in cloacal location repositions external genitalia in amniote evolution. Nature. 2014;516: 391–394. doi:10.1038/nature13819

8. Herrera AM, Cohn MJ. Embryonic origin and compartmental organization of the external genitalia. Sci Rep. 2014;4: 6896. doi:10.1038/srep06896

9. Haraguchi R, Mo R, Hui C, Motoyama J, Makino S, Shiroishi T, et al. Unique functions of Sonic hedgehog signaling during external genitalia development. Development. 2001;128: 4241–4250. doi:10.1242/dev.128.21.4241

10. Perriton CL, Powles N, Chiang C, Maconochie MK, Cohn MJ. Sonic hedgehog Signaling from the Urethral Epithelium Controls External Genital Development. Developmental Biology. 2002;247: 26–46. doi:10.1006/dbio.2002.0668

11. Cohn MJ. Development of the external genitalia: Conserved and divergent mechanisms of appendage patterning. Developmental Dynamics. 2011;240: 1108–1115. doi:10.1002/dvdy.22631

12. Suzuki K, Adachi Y, Numata T, Nakada S, Yanagita M, Nakagata N, et al. Reduced BMP Signaling Results in Hindlimb Fusion with Lethal Pelvic/Urogenital Organ Aplasia: A New Mouse Model of Sirenomelia. Bellusci S, editor. PLoS ONE. 2012;7: e43453. doi:10.1371/journal.pone.0043453

13. Gibson-Brown JJ, Agulnik SI, Chapman DL, Alexiou M, Garvey N, Lee SM, et al. Evidence of a role for T-☐ genes in the evolution of limb morphogenesis and the specification of forelimb/hindlimb identity. Mechanisms of Development. 1996;56: 93–101. doi:10.1016/0925-4773(96)00514-X

14. Rallis C, Bruneau BG, Del Buono J, Seidman CE, Seidman JG, Nissim S, et al. Tbx5 is required for forelimb bud formation and continued outgrowth. Development. 2003;130: 2741–2751. doi:10.1242/dev.00473

15. Lorenzi V, Icoresi-Mazzeo C, Cassie C, Yayon N, Ruiz-Morales ER, Sancho-Serra C, et al. Spatiotemporal cellular map of the developing human reproductive tract. Nature. 2025 [cited 19 Dec 2025]. doi:10.1038/s41586-025-09875-2

16. Matsubara Y, Hirasawa T, Egawa S, Hattori A, Suganuma T, Kohara Y, et al. Anatomical integration of the sacral–hindlimb unit coordinated by GDF11 underlies variation in hindlimb positioning in tetrapods. Nat Ecol Evol. 2017;1: 1392–1399. doi:10.1038/s41559-017-0247-y

17. McPherron AC, Lawler AM, Lee S-J. Regulation of anterior/posterior patterning of the axial skeleton by growth/differentiation factor 11. Nat Genet. 1999;22: 260–264. doi:10.1038/10320

18. Aires R, De Lemos L, Nóvoa A, Jurberg AD, Mascrez B, Duboule D, et al. Tail Bud Progenitor Activity Relies on a Network Comprising Gdf11, Lin28, and Hox13 Genes. Developmental Cell. 2019;48: 383–395.e8. doi:10.1016/j.devcel.2018.12.004

19. Lakshmi G, Abraham J, Benjamin G. Sirenomelia - Revisited. Natl J Clin Anat. 2018;7: 47. doi:10.4103/2277-4025.297648

20. Boulas MM. Recognition of Caudal Regression Syndrome. Advances in Neonatal Care. 2009;9: 61–69. doi:10.1097/ANC.0b013e31819de44f

21. Solomon BD, Raam MS, Pineda Alvarez DE. Analysis of genitourinary anomalies in patients with VACTERL (Vertebral anomalies, Anal atresia, Cardiac malformations, Tracheo Esophageal fistula, Renal anomalies, Limb abnormalities) association. Congenital Anomalies. 2011;51: 87–91. doi:10.1111/j.1741-4520.2010.00303.x

22. Loh KM, Chen A, Koh PW, Deng TZ, Sinha R, Tsai JM, et al. Mapping the Pairwise Choices Leading from Pluripotency to Human Bone, Heart, and Other Mesoderm Cell Types. Cell. 2016;166: 451–467. doi:10.1016/j.cell.2016.06.011

23. Smith CA, Humphreys PA, Naven MA, Woods S, Mancini FE, O’Flaherty J, et al. Directed differentiation of hPSCs through a simplified lateral plate mesoderm protocol for generation of articular cartilage progenitors. Papaccio G, editor. PLoS ONE. 2023;18: e0280024. doi:10.1371/journal.pone.0280024

24. Hofbauer P, Jahnel SM, Papai N, Giesshammer M, Deyett A, Schmidt C, et al. Cardioids reveal self-organizing principles of human cardiogenesis. Cell. 2021;184: 3299–3317.e22. doi:10.1016/j.cell.2021.04.034

25. Zhu J, Zhong X, He H, Cao J, Zhou Z, Dong J, et al. Generation of human expandable limb-bud-like progenitors via chemically induced dedifferentiation. Cell Stem Cell. 2024;31: 1732–1740.e6. doi:10.1016/j.stem.2024.10.001

26. Yamada D, Nakamura M, Takao T, Takihira S, Yoshida A, Kawai S, et al. Induction and expansion of human PRRX1+ limb-bud-like mesenchymal cells from pluripotent stem cells. Nat Biomed Eng. 2021;5: 926–940. doi:10.1038/s41551-021-00778-x

27. Wymeersch FJ, Huang Y, Blin G, Cambray N, Wilkie R, Wong FC, et al. Position-dependent plasticity of distinct progenitor types in the primitive streak. Bronner ME, editor. eLife. 2016;5: e10042. doi:10.7554/eLife.10042

28. Kinder SJ, Tsang TE, Quinlan GA, Hadjantonakis AK, Nagy A, Tam PP. The orderly allocation of mesodermal cells to the extraembryonic structures and the anteroposterior axis during gastrulation of the mouse embryo. Development. 1999;126: 4691–4701. doi:10.1242/dev.126.21.4691

29. Cambray N, Wilson V. Two distinct sources for a population of maturing axial progenitors. Development. 2007;134: 2829–2840. doi:10.1242/dev.02877

30. Wymeersch FJ, Skylaki S, Huang Y, Watson JA, Economou C, Marek-Johnston C, et al. Transcriptionally dynamic progenitor populations organised around a stable niche drive axial patterning. Development. 2019; dev.168161. doi:10.1242/dev.168161

31. Row RH, Pegg A, Kinney BA, Farr GH 3rd, Maves L, Lowell S, et al. BMP and FGF signaling interact to pattern mesoderm by controlling basic helix-loop-helix transcription factor activity. Yelon D, editor. eLife. 2018;7: e31018. doi:10.7554/eLife.31018

32. Hayashi S, Suzuki H, Takemoto T. The nephric mesenchyme lineage of intermediate mesoderm is derived from Tbx6-expressing derivatives of neuro-mesodermal progenitors via BMP-dependent Osr1 function. Developmental Biology. 2021;478: 155–162. doi:10.1016/j.ydbio.2021.07.006

33. Zhang JZ, Termglinchan V, Shao N-Y, Itzhaki I, Liu C, Ma N, et al. A Human iPSC Double-Reporter System Enables Purification of Cardiac Lineage Subpopulations with Distinct Function and Drug Response Profiles. Cell Stem Cell. 2019;24: 802–811.e5. doi:10.1016/j.stem.2019.02.015

34. Frith TJ, Granata I, Wind M, Stout E, Thompson O, Neumann K, et al. Human axial progenitors generate trunk neural crest cells in vitro. eLife. 2018;7: e35786. doi:10.7554/eLife.35786

35. Rojas A, De Val S, Heidt AB, Xu S-M, Bristow J, Black BL. *Gata4* expression in lateral mesoderm is downstream of BMP4 and is activated directly by Forkhead and GATA transcription factors through a distal enhancer element. Development. 2005;132: 3405–3417. doi:10.1242/dev.01913

36. Newton AH, Smith CA. Resolving the mechanisms underlying epithelial to mesenchymal transition of the lateral plate mesoderm. Genesis. 2024;62: e23531. doi:10.1002/dvg.23531

37. Thomas T, Yamagishi H, Overbeek PA, Olson EN, Srivastava D. The bHLH Factors, dHAND and eHAND, Specify Pulmonary and Systemic Cardiac Ventricles Independent of Left–Right Sidedness. Developmental Biology. 1998;196: 228–236. doi:10.1006/dbio.1998.8849

38. Lynch AT, Phillips N, Douglas M, Dorgnach M, Lin I-H, Adamson AD, et al. HAND1 level controls the specification of multipotent cardiac and extraembryonic progenitors from human pluripotent stem cells. EMBO J. 2025;44: 8. doi:10.1038/s44318-025-00409-0

39. Gattiglio M, Protzek M, Schröter C. Population-level antagonism between FGF and BMP signaling steers mesoderm differentiation in embryonic stem cells. Biology Open. 2023;12: bio059941. doi:10.1242/bio.059941

40. Barnes RM, Firulli BA, Conway SJ, Vincentz JW, Firulli AB. Analysis of the Hand1 cell lineage reveals novel contributions to cardiovascular, neural crest, extra embryonic, and lateral mesoderm derivatives. Developmental Dynamics. 2010;239: 3086–3097. doi:10.1002/dvdy.22428

41. Fazilaty H, Rago L, Kass Youssef K, Ocaña OH, Garcia-Asencio F, Arcas A, et al. A gene regulatory network to control EMT programs in development and disease. Nat Commun. 2019;10: 5115. doi:10.1038/s41467-019-13091-8

42. Marcil A, Dumontier É, Chamberland M, Camper SA, Drouin J. *Pitx1* and *Pitx2* are required for development of hindlimb buds. Development. 2003;130: 45–55. doi:10.1242/dev.00192

43. Lanctôt C, Lamolet B, Drouin J. The *bicoid* -related homeoprotein *Ptx1* defines the most anterior domain of the embryo and differentiates posterior from anterior lateral mesoderm. Development. 1997;124: 2807–2817. doi:10.1242/dev.124.14.2807

44. Foerst-Potts L, Sadler TW. Disruption ofMsx-1 andMsx-2 reveals roles for these genes in craniofacial, eye, and axial development. Dev Dyn. 1997;209: 70–84. doi:10.1002/(SICI)1097-0177(199705)209:1<70::AID-AJA7>3.0.CO;2-U

45. Rouco R, Bompadre O, Rauseo A, Fazio O, Peraldi R, Thorel F, et al. Cell-specific alterations in Pitx1 regulatory landscape activation caused by the loss of a single enhancer. Nat Commun. 2021;12: 7235. doi:10.1038/s41467-021-27492-1

46. Armfield BA, Cohn MJ. Single cell transcriptomic analysis of external genitalia reveals complex and sexually dimorphic cell populations in the early genital tubercle. Developmental Biology. 2021;477: 145–154. doi:10.1016/j.ydbio.2021.05.014

47. Houweling AC, Dildrop R, Peters T, Mummenhoff J, Moorman AFM, Rüther U, et al. Gene and cluster-specific expression of the Iroquois family members during mouse development. Mechanisms of Development. 2001;107: 169–174. doi:10.1016/S0925-4773(01)00451-8

48. Cobb J, Duboule D. Comparative analysis of genes downstream of the Hoxd cluster in developing digits and external genitalia. Development. 2005;132: 3055–3067. doi:10.1242/dev.01885

49. Sefton M, Sánchez S, Nieto MA. Conserved and divergent roles for members of the *Snail* family of transcription factors in the chick and mouse embryo. Development. 1998;125: 3111–3121. doi:10.1242/dev.125.16.3111

50. Nieto MA, Sargent MG, Wilkinson DG, Cooke J. Control of Cell Behavior During Vertebrate Development by *Slug* , a Zinc Finger Gene. Science. 1994;264: 835–839. doi:10.1126/science.7513443

51. Chapman DL, Garvey N, Hancock S, Alexiou M, Agulnik SI, Gibson-Brown JJ, et al. Expression of the T-box family genes,Tbx1-Tbx5, during early mouse development. Dev Dyn. 1996;206: 379–390. doi:10.1002/(SICI)1097-0177(199608)206:4<379::AID-AJA4>3.0.CO;2-F

52. Inman KE, Downs KM. Localization of Brachyury (T) in embryonic and extraembryonic tissues during mouse gastrulation. Gene Expression Patterns. 2006;6: 783–793. doi:10.1016/j.modgep.2006.01.010

53. Inman KE, Downs KM. Brachyury is required for elongation and vasculogenesis in the murine allantois. Development. 2006;133: 2947–2959. doi:10.1242/dev.02454

54. Cheifetz S, Bellón T, Calés C, Vera S, Bernabeu C, Massagué J, et al. Endoglin is a component of the transforming growth factor-beta receptor system in human endothelial cells. J Biol Chem. 1992;267: 19027–19030.

55. Lee C, Kim M-J, Kumar A, Lee H-W, Yang Y, Kim Y. Vascular endothelial growth factor signaling in health and disease: from molecular mechanisms to therapeutic perspectives. Sig Transduct Target Ther. 2025;10: 170. doi:10.1038/s41392-025-02249-0

56. Richardson MR, Robbins EP, Vemula S, Critser PJ, Whittington C, Voytik-Harbin SL, et al. Angiopoietin-like protein 2 regulates endothelial colony forming cell vasculogenesis. Angiogenesis. 2014;17: 675–683. doi:10.1007/s10456-014-9423-8

57. Prummel KD, Nieuwenhuize S, Mosimann C. The lateral plate mesoderm. Development. 2020;147: dev175059. doi:10.1242/dev.175059

58. Dollé P, Izpisúa-Belmonte JC, Brown JM, Tickle C, Duboule D. HOX-4 genes and the morphogenesis of mammalian genitalia. Genes Dev. 1991;5: 1767–1776. doi:10.1101/gad.5.10.1767

59. Zeltser L, Desplan C, Heintz N. *Hoxb-13* : a new Hox gene in a distant region of the HOXB cluster maintains colinearity. Development. 1996;122: 2475–2484. doi:10.1242/dev.122.8.2475

60. Lopez-Yrigoyen M, Fidanza A, Cassetta L, Axton RA, Taylor AH, Meseguer-Ripolles J, et al. A human iPSC line capable of differentiating into functional macrophages expressing ZsGreen: a tool for the study and *in vivo* tracking of therapeutic cells. Phil Trans R Soc B. 2018;373: 20170219. doi:10.1098/rstb.2017.0219

61. Thomson JA, Itskovitz-Eldor J, Shapiro SS, Waknitz MA, Swiergiel JJ, Marshall VS, et al. Embryonic stem cell lines derived from human blastocysts. Science. 1998;282: 1145–1147. doi:10.1126/science.282.5391.1145

62. Deimling SJ, Drysdale TA. Retinoic acid regulates anterior–posterior patterning within the lateral plate mesoderm of Xenopus. Mechanisms of Development. 2009;126: 913–923. doi:10.1016/j.mod.2009.07.001

63. Bernheim S, Meilhac SM. Mesoderm patterning by a dynamic gradient of retinoic acid signalling. Phil Trans R Soc B. 2020;375: 20190556. doi:10.1098/rstb.2019.0556

64. Liu L, Suzuki K, Nakagata N, Mihara K, Matsumaru D, Ogino Y, et al. Retinoic Acid Signaling Regulates Sonic Hedgehog and Bone Morphogenetic Protein Signalings During Genital Tubercle Development. Birth Defects Research Pt B. 2012;95: 79–88. doi:10.1002/bdrb.20344

65. Millauer B, Wizigmann-Voos S, Schnürch H, Martinez R, Møller NPH, Risau W, et al. High affinity VEGF binding and developmental expression suggest Flk-1 as a major regulator of vasculogenesis and angiogenesis. Cell. 1993;72: 835–846. doi:10.1016/0092-8674(93)90573-9

66. Yamaguchi TP, Dumont DJ, Conlon RA, Breitman ML, Rossant J. *flk-1* , an *flt* -related receptor tyrosine kinase is an early marker for endothelial cell precursors. Development. 1993;118: 489–498. doi:10.1242/dev.118.2.489

67. Arora R, Metzger RJ, Papaioannou VE. Multiple Roles and Interactions of Tbx4 and Tbx5 in Development of the Respiratory System. Barsh GS, editor. PLoS Genet. 2012;8: e1002866. doi:10.1371/journal.pgen.1002866

68. Humphreys PEA, Woods S, Bates N, Rooney KM, Mancini FE, Barclay C, et al. Optogenetic manipulation of BMP signaling to drive chondrogenic differentiation of hPSCs. Cell Reports. 2023;42: 113502. doi:10.1016/j.celrep.2023.113502

69. Douglas NC, Heng K, Sauer MV, Papaioannou VE. Dynamic expression of Tbx2 subfamily genes in development of the mouse reproductive system. Developmental Dynamics. 2012;241: 365–375. doi:10.1002/dvdy.23710

70. Mills AA, Zheng B, Wang X-J, Vogel H, Roop DR, Bradley A. p63 is a p53 homologue required for limb and epidermal morphogenesis. Nature. 1999;398: 708–713. doi:10.1038/19531

71. Tanaka K, Matsumaru D, Suzuki K, Yamada G, Miyagawa S. The role of p63 in embryonic external genitalia outgrowth in mice. Dev Growth Differ. 2023;65: 132–140. doi:10.1111/dgd.12840

72. Haro E, Delgado I, Junco M, Yamada Y, Mansouri A, Oberg KC, et al. Sp6 and Sp8 Transcription Factors Control AER Formation and Dorsal-Ventral Patterning in Limb Development. Lewandoski M, editor. PLoS Genet. 2014;10: e1004468. doi:10.1371/journal.pgen.1004468

73. Lin C, Yin Y, Bell SM, Veith GM, Chen H, Huh S-H, et al. Delineating a Conserved Genetic Cassette Promoting Outgrowth of Body Appendages. Beier DR, editor. PLoS Genet. 2013;9: e1003231. doi:10.1371/journal.pgen.1003231

74. Ferrari D, Sumoy L, Gannon J, Sun H, Brown AMC, Upholt WB, et al. The expression pattern of the Distal-less homeo☐-containing gene Dlx-5 in the developing chick limb bud suggests its involvement in apical ectodermal ridge activity, pattern formation, and cartilage differentiation. Mechanisms of Development. 1995;52: 257–264. doi:10.1016/0925-4773(95)98113-O

75. Robledo RF, Rajan L, Li X, Lufkin T. The *Dlx5* and *Dlx6* homeobox genes are essential for craniofacial, axial, and appendicular skeletal development. Genes Dev. 2002;16: 1089–1101. doi:10.1101/gad.988402

76. Dealy CN, Roth A, Ferrari D, Brown AMC, Kosher RA. Wnt-5a and Wnt-7a are expressed in the developing chick limb bud in a manner suggesting roles in pattern formation along the proximodistal and dorsoventral axes. Mechanisms of Development. 1993;43: 175–186. doi:10.1016/0925-4773(93)90034-U

77. Yamaguchi TP, Bradley A, McMahon AP, Jones S. A *Wnt5a* pathway underlies outgrowth of multiple structures in the vertebrate embryo. Development. 1999;126: 1211–1223. doi:10.1242/dev.126.6.1211

78. Duboc V, Sulaiman F, Feneck E, Kucharska A, Bell D, Holder-Espinasse M, et al. *Tbx4* function during hindlimb development reveals a novel mechanism to explain the origins of proximal limb defects. Development. 2021; dev.199580. doi:10.1242/dev.199580

79. Hasson P, DeLaurier A, Bennett M, Grigorieva E, Naiche LA, Papaioannou VE, et al. Tbx4 and Tbx5 Acting in Connective Tissue Are Required for Limb Muscle and Tendon Patterning. Developmental Cell. 2010;18: 148–156. doi:10.1016/j.devcel.2009.11.013

80. Xie T, Liang J, Liu N, Huan C, Zhang Y, Liu W, et al. Transcription factor TBX4 regulates myofibroblast accumulation and lung fibrosis. Journal of Clinical Investigation. 2016;126: 3063–3079. doi:10.1172/JCI85328

81. Pan J, Li Y, Lin Z, Lan Q, Zhang Y, Chen H, et al. Spatiotemporal transcriptome atlas of human embryos after gastrulation. Nature. 2026;654: 751–761. doi:10.1038/s41586-026-10545-0

82. Pan J, Lin Z, Zhang G, Huang H, Ding G. Charting human organogenesis across the Carnegie stages from a whole-embryo perspective. GigaScience. 2026;15: giag063. doi:10.1093/gigascience/giag063

83. Cao J, Spielmann M, Qiu X, Huang X, Ibrahim DM, Hill AJ, et al. The single-cell transcriptional landscape of mammalian organogenesis. Nature. 2019;566: 496–502. doi:10.1038/s41586-019-0969-x

84. Gipson GR, Goebel EJ, Hart KN, Kappes EC, Kattamuri C, McCoy JC, et al. Structural perspective of BMP ligands and signaling. Bone. 2020;140: 115549. doi:10.1016/j.bone.2020.115549

85. Zhang Y-H, Cheng F, Du X-T, Gao J-L, Xiao X-L, Li N, et al. GDF11/BMP11 activates both smad1/5/8 and smad2/3 signals but shows no significant effect on proliferation and migration of human umbilical vein endothelial cells. Oncotarget. 2016;7: 12063–12074. doi:10.18632/oncotarget.7642

86. Auth JM, Eizenberg-Magar I, Erez O, Zachar O, Shmueli MD, Zigdon I, et al. Asymmetric crosstalk between the BMP and TGFβ pathways resolves signaling ambiguity. bioRxiv. 2026; 2026.04.20.719570. doi:10.64898/2026.04.20.719570

87. Tonegawa A, Funayama N, Ueno N, Takahashi Y. Mesodermal subdivision along the mediolateral axis in chicken controlled by different concentrations of BMP-4. Development. 1997;124: 1975–1984. doi:10.1242/dev.124.10.1975

88. Tonegawa A, Takahashi Y. Somitogenesis controlled by Noggin. Dev Biol. 1998;202: 172–182. doi:10.1006/dbio.1998.8895

89. Leung AW, Murdoch B, Salem AF, Prasad MS, Gomez GA, García-Castro MI. WNT/β-catenin signaling mediates human neural crest induction via a pre-neural border intermediate. Development. 2016;143: 398–410. doi:10.1242/dev.130849

90. Nakanoh S, Sham K, Ghimire S, Mohorianu I, Rayon T, Vallier L. Human surface ectoderm and amniotic ectoderm are sequentially specified according to cellular density. Sci Adv. 2024;10: eadh7748. doi:10.1126/sciadv.adh7748

91. Skoufa E, Zhong J, Hu K, Kahre O, Tsissios G, Carrau L, et al. Specialized signaling centers direct cell fate and spatial organization in a mesodermal organoid model. Sci Adv. 2025;11: eady7682. doi:10.1126/sciadv.ady7682

92. Mori S, Sakakura E, Tsunekawa Y, Hagiwara M, Suzuki T, Eiraku M. Self-organized formation of developing appendages from murine pluripotent stem cells. Nat Commun. 2019;10: 3802. doi:10.1038/s41467-019-11702-y

93. The International Stem Cell Initiative. Screening ethnically diverse human embryonic stem cells identifies a chromosome 20 minimal amplicon conferring growth advantage. Nat Biotechnol. 2011;29: 1132–1144. doi:10.1038/nbt.2051

94. Schindelin J, Arganda-Carreras I, Frise E, Kaynig V, Longair M, Pietzsch T, et al. Fiji: an open-source platform for biological-image analysis. Nat Methods. 2012;9: 676–682. doi:10.1038/nmeth.2019

95. Stirling DR, Swain-Bowden MJ, Lucas AM, Carpenter AE, Cimini BA, Goodman A. CellProfiler 4: improvements in speed, utility and usability. BMC Bioinformatics. 2021;22: 433. doi:10.1186/s12859-021-04344-9

96. Mortazavi A, Williams BA, McCue K, Schaeffer L, Wold B. Mapping and quantifying mammalian transcriptomes by RNA-Seq. Nat Methods. 2008;5: 621–628. doi:10.1038/nmeth.1226

97. Liao Y, Smyth GK, Shi W. featureCounts: an efficient general purpose program for assigning sequence reads to genomic features. Bioinformatics. 2014;30: 923–930. doi:10.1093/bioinformatics/btt656

98. Love MI, Huber W, Anders S. Moderated estimation of fold change and dispersion for RNA-seq data with DESeq2. Genome Biol. 2014;15: 550. doi:10.1186/s13059-014-0550-8

99. Yu G, Wang L-G, Han Y, He Q-Y. clusterProfiler: an R package for comparing biological themes among gene clusters. OMICS. 2012;16: 284–287. doi:10.1089/omi.2011.0118

100. Stuart T, Butler A, Hoffman P, Hafemeister C, Papalexi E, Mauck WM, et al. Comprehensive Integration of Single-Cell Data. Cell. 2019;177: 1888–1902.e21. doi:10.1016/j.cell.2019.05.031

101. McGinnis CS, Murrow LM, Gartner ZJ. DoubletFinder: Doublet Detection in Single-Cell RNA Sequencing Data Using Artificial Nearest Neighbors. Cell Systems. 2019;8: 329–337.e4. doi:10.1016/j.cels.2019.03.003

102. Samuel Marsh, Maëlle Salmon, Paul Hoffman, kew24, Mustafa Samet Pir. samuel-marsh/scCustomize: Release 3.2.4. Zenodo; 2025. doi:10.5281/ZENODO.5706430

103. Trapnell C, Cacchiarelli D, Grimsby J, Pokharel P, Li S, Morse M, et al. The dynamics and regulators of cell fate decisions are revealed by pseudotemporal ordering of single cells. Nat Biotechnol. 2014;32: 381–386. doi:10.1038/nbt.2859

104. Patikas N, Yao H, Madhu R, Raychaudhuri S, Hemberg M, Korsunsky I. Integration of large, complex single-cell datasets with Harmony2. Bioinformatics; 2026. doi:10.64898/2026.03.16.711825

105. Choi HMT, Schwarzkopf M, Fornace ME, Acharya A, Artavanis G, Stegmaier J, et al. Third-generation *in situ* hybridization chain reaction: multiplexed, quantitative, sensitive, versatile, robust. Development. 2018;145: dev165753. doi:10.1242/dev.165753

106. Zeng B, Liu Z, Lu Y, Zhong S, Qin S, Huang L, et al. The single-cell and spatial transcriptional landscape of human gastrulation and early brain development. Cell Stem Cell. 2023;30: 851–866.e7. doi:10.1016/j.stem.2023.04.016

