## Supplementary material for "Generation of human hindlimb/genital tubercle progenitors from pluripotent stem cells": Suppl Figures

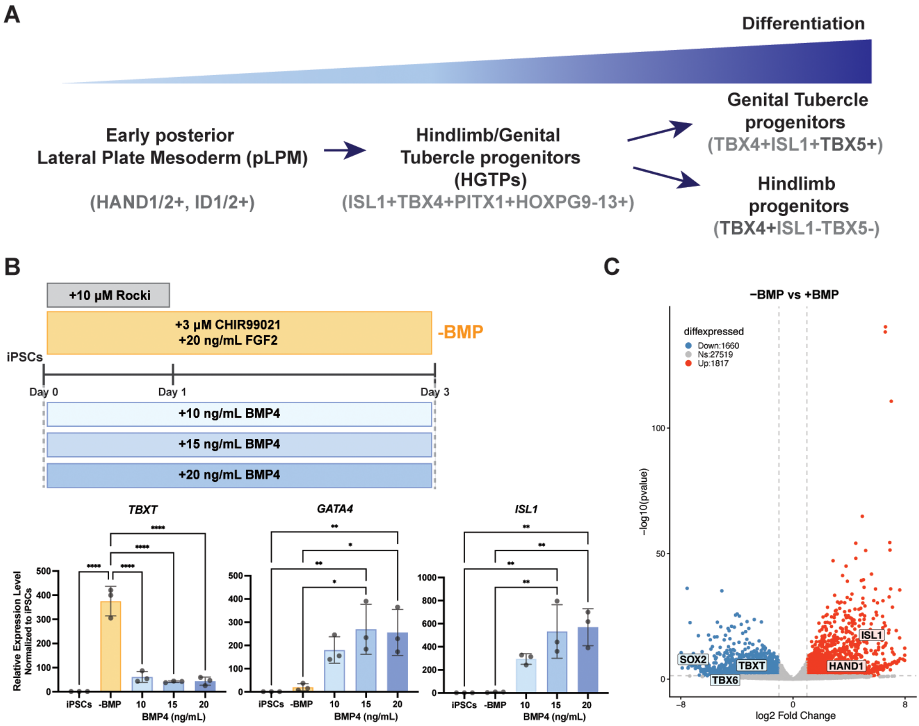
A
Differentiation
Early posterior
Lateral Plate Mesoderm (pLPM)
(HAND1/2+, ID1/2+)
Hindlimb/Genital
Tubercle progenitors
(HGTPs)
(ISL1+TBX4+PITX1+HOXPG9-13+)
Genital Tubercle
progenitors
(TBX4+ISL1+TBX5+)
Hindlimb
progenitors
(TBX4+ISL1-TBX5-)
B
+10 μM Rocki
+3 μM CHIR99021
+20 ng/mL FGF2
-BMP
iPSCs
Day 0
Day 1
Day 3
+10 ng/mL BMP4
+15 ng/mL BMP4
+20 ng/mL BMP4
TBXT
GATA4
ISL1
Relative Expression Level
Normalized to iPSCs
iPSCs
-BMP
BMP4 (ng/mL)
C
−BMP vs +BMP
diffexpressed
Down:1660
Ns:27519
Up:1817
−log10(pvalue)
log2 Fold Change
SOX2
TBXT
TBX6
ISL1
HAND1

S1

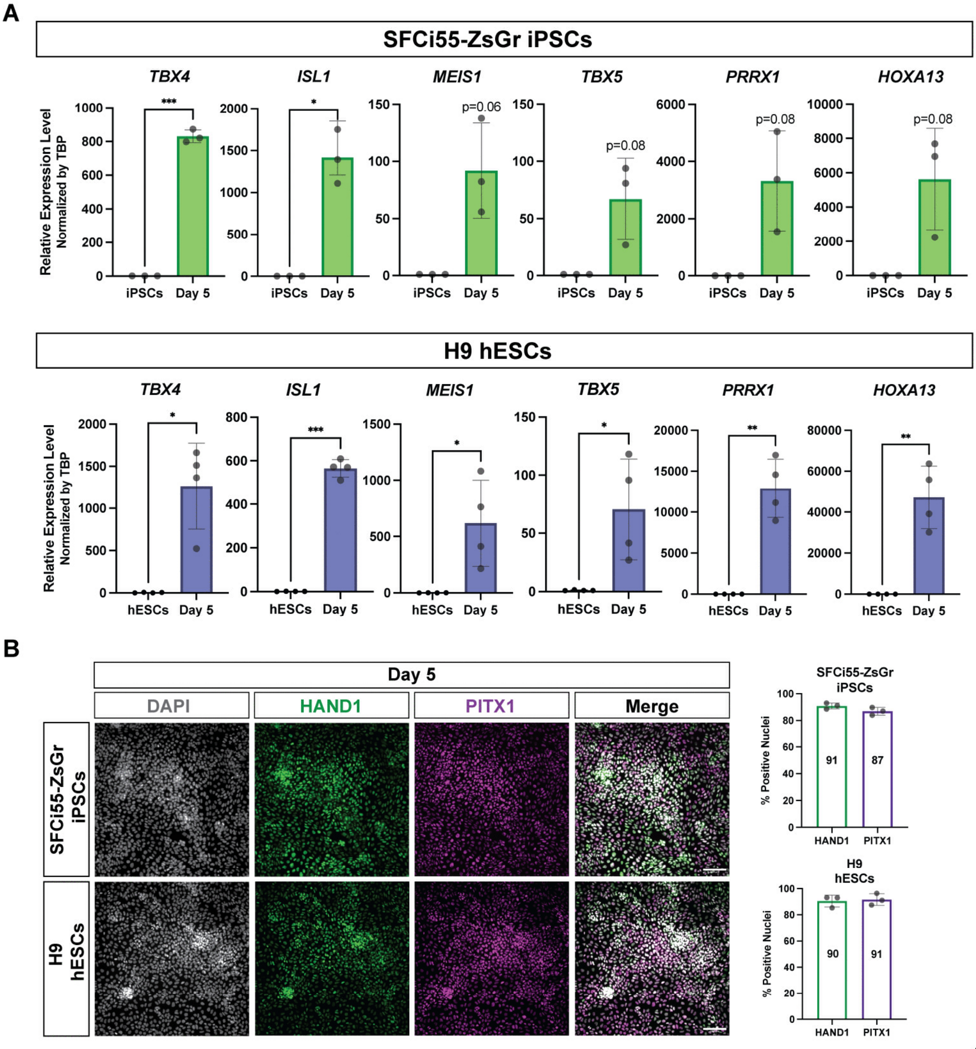
A
SFCi55-ZsGr iPSCs
TBX4
ISL1
MEIS1
TBX5
PRRX1
HOXA13
Relative Expression Level
Normalized by TBP
***
*
p=0.06
p=0.08
p=0.08
p=0.08
iPSCs
Day 5
H9 hESCs
TBX4
ISL1
MEIS1
TBX5
PRRX1
HOXA13
Relative Expression Level
Normalized by TBP
*
***
*
*
**
**
hESCs
Day 5
B
Day 5
DAPI
HAND1
PITX1
Merge
SFCi55-ZsGr iPSCs
H9 hESCs
SFCi55-ZsGr iPSCs
% Positive Nuclei
91
87
HAND1
PITX1
H9 hESCs
% Positive Nuclei
90
91
HAND1
PITX1

S2

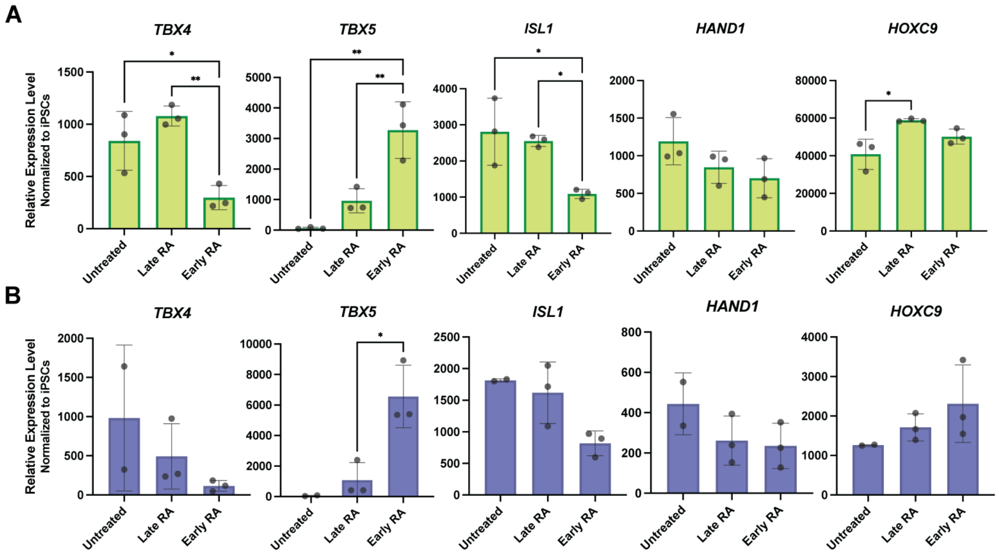
A
TBX4
TBX5
ISL1
HAND1
HOXC9
Relative Expression Level
Normalized to iPSCs
Untreated
Late RA
Early RA
B
TBX4
TBX5
ISL1
HAND1
HOXC9
Relative Expression Level
Normalized to iPSCs
Untreated
Late RA
Early RA

S3

**A**

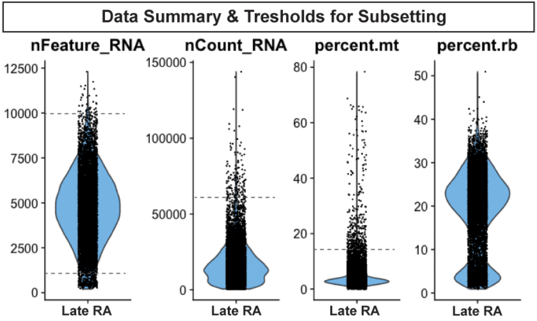


**B**

| | Cell Counts | |
|---|---|---|
| | Before Filtering | After Filtering |
| Late RA | 9122 | 8292 |

**C**

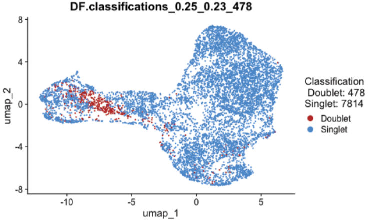


**S4**

S5

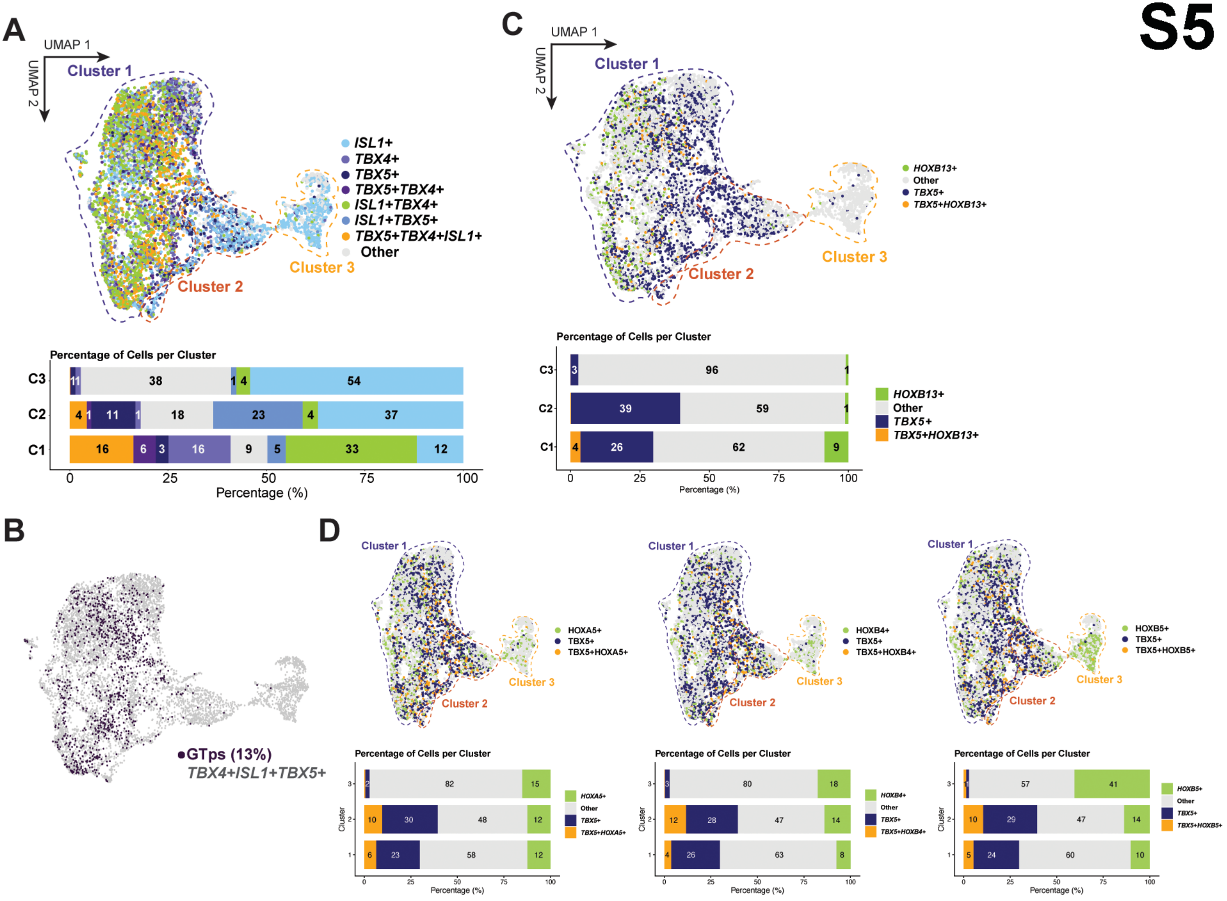
A
UMAP 1
UMAP 2
Cluster 1
Cluster 2
Cluster 3
ISL1+
TBX4+
TBX5+
TBX5+TBX4+
ISL1+TBX4+
ISL1+TBX5+
TBX5+TBX4+ISL1+
Other
Percentage of Cells per Cluster
C3 1 1 38 1 4 54
C2 4 1 11 1 18 23 4 37
C1 16 6 3 16 9 5 33 12
0 25 50 75 100
Percentage (%)
B
●GTps (13%)
TBX4+ISL1+TBX5+
C
UMAP 1
UMAP 2
Cluster 1
Cluster 2
Cluster 3
HOXB13+
Other
TBX5+
TBX5+HOXB13+
Percentage of Cells per Cluster
C3 3 96 1
C2 39 59 1
C1 4 26 62 9
0 25 50 75 100
Percentage (%)
HOXB13+
Other
TBX5+
TBX5+HOXB13+
D
Cluster 1
Cluster 2
Cluster 3
HOXA5+
TBX5+
TBX5+HOXA5+
Percentage of Cells per Cluster
Cluster
3 2 82 15
2 10 30 48 12
1 6 23 58 12
0 25 50 75 100
Percentage (%)
HOXA5+
Other
TBX5+
TBX5+HOXA5+
Cluster 1
Cluster 2
Cluster 3
HOXB4+
TBX5+
TBX5+HOXB4+
Percentage of Cells per Cluster
Cluster
3 3 80 18
2 12 28 47 14
1 4 26 63 8
0 25 50 75 100
Percentage (%)
HOXB4+
Other
TBX5+
TBX5+HOXB4+
Cluster 1
Cluster 2
Cluster 3
HOXB5+
TBX5+
TBX5+HOXB5+
Percentage of Cells per Cluster
Cluster
3 1 57 41
2 10 29 47 14
1 5 24 60 10
0 25 50 75 100
Percentage (%)
HOXB5+
Other
TBX5+
TBX5+HOXB5+

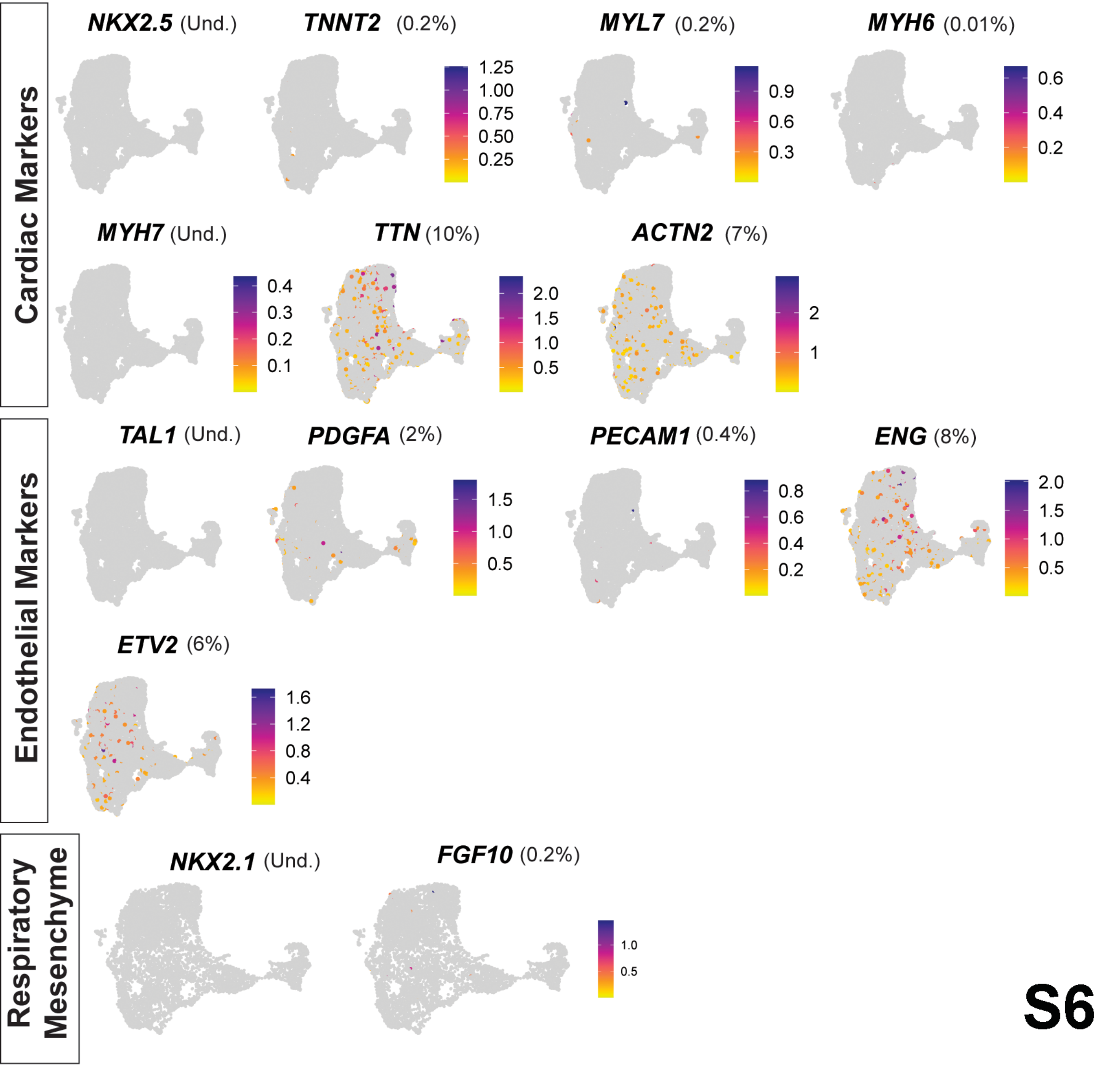
Cardiac Markers
NKX2.5 (Und.)
TNNT2 (0.2%)
1.25
1.00
0.75
0.50
0.25
MYL7 (0.2%)
0.9
0.6
0.3
MYH6 (0.01%)
0.6
0.4
0.2
MYH7 (Und.)
0.4
0.3
0.2
0.1
TTN (10%)
2.0
1.5
1.0
0.5
ACTN2 (7%)
2
1
Endothelial Markers
TAL1 (Und.)
PDGFA (2%)
1.5
1.0
0.5
PECAM1 (0.4%)
0.8
0.6
0.4
0.2
ENG (8%)
2.0
1.5
1.0
0.5
ETV2 (6%)
1.6
1.2
0.8
0.4
Respiratory Mesenchyme
NKX2.1 (Und.)
FGF10 (0.2%)
1.0
0.5

**S6**

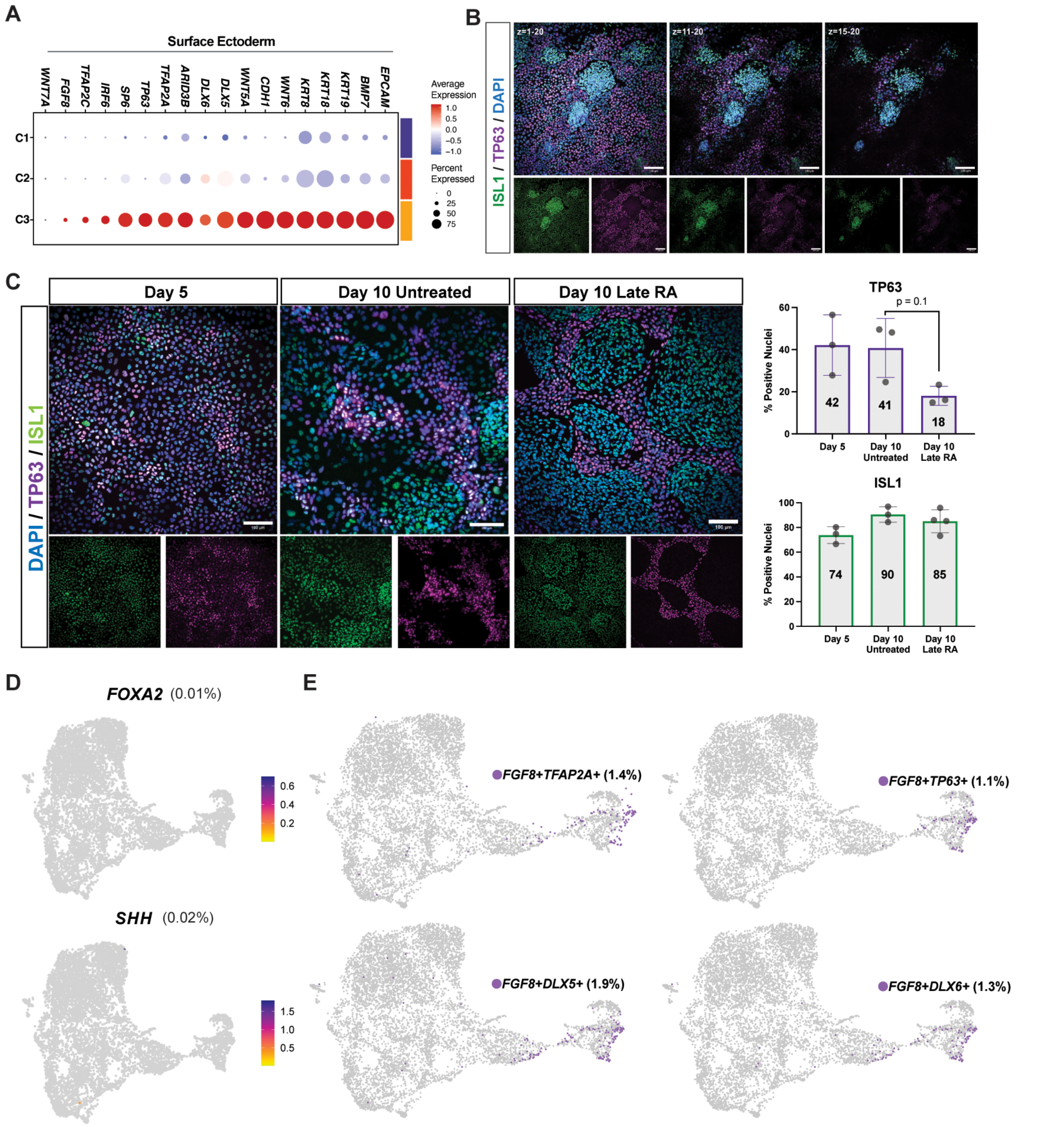

A
Surface Ectoderm
WNT7A
FGF8
TFAP2C
IRF6
SP6
TP63
TFAP2A
ARID3B
DLX6
DLX5
WNT5A
CDH1
WNT6
KRT8
KRT18
KRT19
BMP7
EPCAM
C1
C2
C3
Average Expression
1.0
0.5
0.0
−0.5
−1.0
Percent Expressed
0
25
50
75
B
ISL1 / TP63 / DAPI
z=1-20
z=11-20
z=15-20
C
Day 5
Day 10 Untreated
Day 10 Late RA
DAPI / TP63 / ISL1
TP63
p = 0.1
% Positive Nuclei
42
41
18
Day 5
Day 10 Untreated
Day 10 Late RA
ISL1
% Positive Nuclei
74
90
85
Day 5
Day 10 Untreated
Day 10 Late RA
D
FOXA2 (0.01%)
0.6
0.4
0.2
SHH (0.02%)
1.5
1.0
0.5
E
FGF8+TFAP2A+ (1.4%)
FGF8+TP63+ (1.1%)
FGF8+DLX5+ (1.9%)
FGF8+DLX6+ (1.3%)


S7

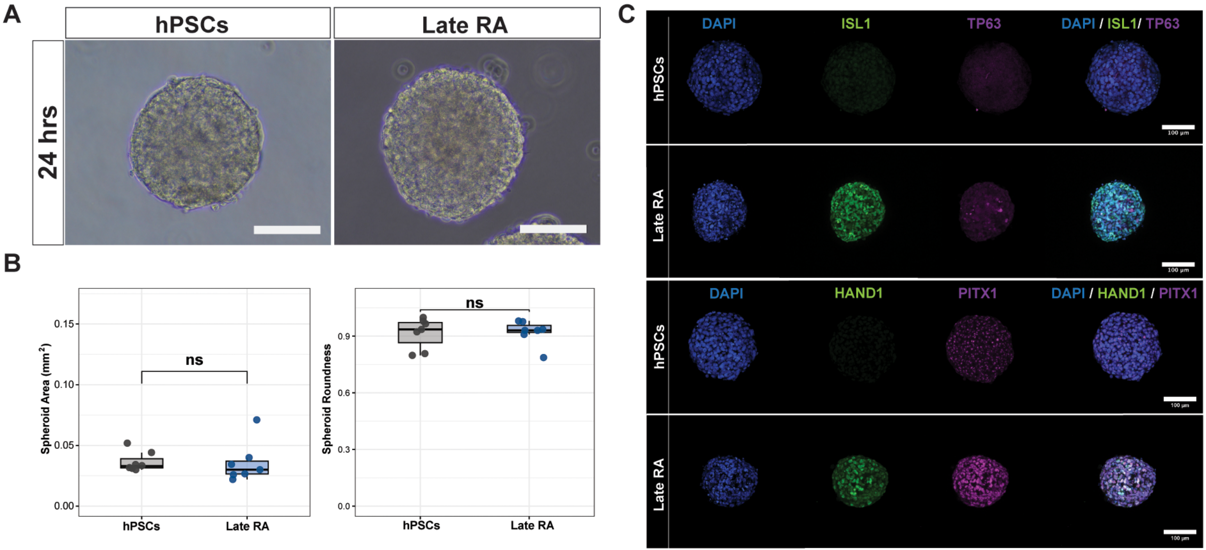
A
hPSCs
Late RA
24 hrs
B
Spheroid Area (mm²)
0.15
0.10
0.05
0.00
ns
hPSCs
Late RA
Spheroid Roundness
0.9
0.6
0.3
0.0
ns
hPSCs
Late RA
C
DAPI
ISL1
TP63
DAPI / ISL1/ TP63
hPSCs
Late RA
DAPI
HAND1
PITX1
DAPI / HAND1 / PITX1
hPSCs
Late RA
100 µm

S8

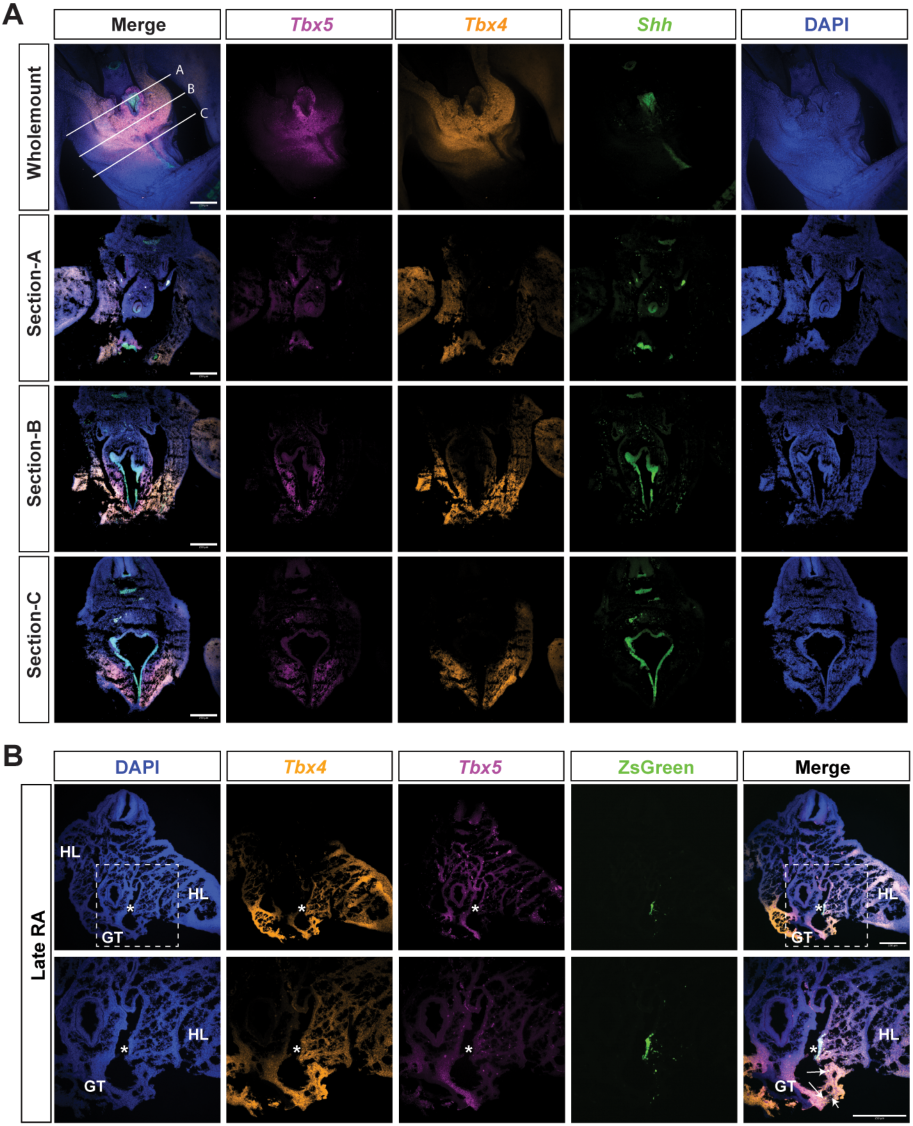
A
Merge
Tbx5
Tbx4
Shh
DAPI
Wholemount
Section-A
Section-B
Section-C
A
B
C
B
DAPI
Tbx4
Tbx5
ZsGreen
Merge
Late RA
HL
GT
*

S9
